# Antagonism between viral infection and innate immunity at the single-cell level

**DOI:** 10.1101/2022.11.18.517110

**Authors:** Frederic Grabowski, Marek Kochańczyk, Zbigniew Korwek, Maciej Czerkies, Wiktor Prus, Tomasz Lipniacki

**Affiliations:** Institute of Fundamental Technological Research, Polish Academy of Sciences, 02-106 Warsaw, Poland

**Author notes:** These authors contributed equally to this work (Frederic Grabowski, Marek Kochańczyk).

**Keywords:** agent-based modeling, immunofluorescence imaging, mutual inhibition, respiratory syncytial virus, virus–host interactions

## Abstract

When infected with a virus, cells may secrete interferons (IFNs) that prompt nearby cells to prepare for upcoming infection. Reciprocally, viral proteins often interfere with IFN synthesis and IFN-induced signaling. We modeled the crosstalk between the propagating virus and the innate immune response using an agent-based stochastic approach. By analyzing immunofluorescence microscopy images we observed that the mutual antagonism between the respiratory syncytial virus (RSV) and infected A549 cells leads to dichotomous responses at the single-cell level and complex spatial patterns of cell signaling states. Our analysis indicates that RSV blocks innate responses at three levels: by inhibition of IRF3 activation, inhibition of IFN synthesis, and inhibition of STAT1/2 activation. In turn, proteins coded by IFN-stimulated (STAT1/2-activated) genes inhibit the synthesis of viral RNA and viral proteins. The striking consequence of these inhibitions is lack of coincidence of viral proteins and IFN expression within single cells. The model enables investigation of the impact of immunostimulatory defective viral particles and signaling network perturbations that could potentially facilitate containment or clearance of the viral infection.

## Introduction

Innate immunity provides the first line of defense against viral infections. It acts primarily by feed-forward signaling from infected cells to not (yet) infected bystander cells: upon recognition of viral genetic material by intracellular receptors, some infected cells synthesize and secrete intercellular messengers, most importantly interferons (IFNs), that prompt nearby cells to enter an antiviral state, rendering them more resistant to secondary infections (Czerkies *et al*, 2018; Patil *et al*, 2015; Rand *et al*, 2012; Schoggins, 2019; Zhao *et al*, 2012). Imminently, with respect to the presence or absence of the virus and the antiviral state, a (partially) infected cell population is spatiotemporally stratified into functionally distinct subpopulations (Drayman *et al*, 2019; Korwek *et al*, 2022; Rand *et al*, 2012; Sen *et al*, 2012).

To invade a host cell and successfully replicate, viruses evolved multiple strategies to evade or counteract cellular mechanisms of the innate immune response (García-Sastre, 2017). Most often, viral proteins, in particular non-structural proteins, attenuate viral RNA sensing and interfere with synthesis of IFNs and IFN-induced STAT signaling. For example, in respiratory viruses, NS1 of influenza A/B virus (IAV/IBV) directly and indirectly inhibits activation of IRF3, a key transcription factor of type I and type III IFNs (Ayllon & García-Sastre, 2015; Hale *et al*, 2010); NS1 and NS2 of the respiratory syncytial virus (RSV) similarly impinge on activation of IRF3 and, by disrupting STAT1 and STAT2 signaling, impede the induction of IFN-stimulated genes (ISGs) (Barik, 2013; Sedeyn *et al*, 2019); a multitude of effectively analogous interactions have been recently discovered in the case of SARS-CoV-2 (Rashid *et al*, 2022; Sa Ribero *et al*, 2020). The mutual virus–host cell interactions amplify heterogeneity of the stratified, infected cell population and modulate the progression of viral infection. The picture can be even more convoluted in the presence of immunostimulatory, replication-incompetent defective viral particles (Vignuzzi & López, 2019).

Understanding the complex interplay between viruses and the innate immune response at the system level requires experimentation at the single-cell resolution combined with data-driven single-cell level computational modeling (Talemi & Höfer, 2018; Van Eyndhoven *et al*, 2021). Thus far, integration of quantitative data on *Ifnb1* (IFNβ gene) and ISGs expression in single cells over time and stochastic modeling allowed to demonstrate that paracrine signaling has a major impact on heterogeneous cell responses to infection (Patil *et al*, 2015; Rand *et* al, 2012; Maier *et al*, 2022). Recently, benefits of interferon expression heterogeneity have been studied within a spatial agent-based model of an infection with a “generic” virus (Gregg *et al*, 2021). A spatial agent-based model with ODE-based intracellular kinetics and stochastic transitions between cell types in a recent work of Aponte-Serrano *et al* (2021) was directly tuned to recapitulate the kinetics of plaque formation by IAV in human bronchial cells. It was demonstrated that a sufficiently fast synthesis and diffusion of (or a prestimulation with) an interferon may entirely arrest plaque growth. The action of IAV NS1 was factored into a reduction of the activity of a viral RNA sensor and an IRF inducer, RIG-I. The employed generative computational approach enabled extending the model by inclusion of multiple types of immune cell types (also those involved in adaptive immunity), several diffusible cytokines, as well as cell attachment and migration (Sego *et al,* 2021, 2022). It was proposed that locally concentrated exposure to a virus is able to elicit a productive infection, whereas uniform exposure to virus is likely ineffective.

Here, we present an agent-based, spatial, stochastic model of infection propagation in a monolayer of cells. The model was developed and calibrated based on an array of our experiments on epithelial cells of respiratory origin (A549 cell line) infected with RSV, the most common pathogen in severe respiratory disease in both young children and the elderly (Crowe, 2014; Falsey *et al*, 2005; Scheltema *et al*, 2017). Upon viral infection, these cells communicate through interferons of type I (IFNβ) and type III (IFNλ) (Czerkies *et al*, 2022). By analyzing results from cell population-level techniques (Western blot, ELISA, dPCR), we delineated the structure and constrained kinetic rates of the regulatory network. By analyzing heterogeneous single-cell responses in immunofluorescence microscopy images, we quantified the degree of antagonism between the virus and host cell. The data demonstrate that epithelial cells fight the virus using STAT-inducible ISGs, which inhibit synthesis of viral RNA and proteins. RSV proteins, in turn, attenuate immune responses by interfering with the activation of IRF3, synthesis of IFNs, and phosphorylation of STAT1/2. These reciprocal, antagonistic interactions give rise to a transient switch-like behavior: an infected cell is unlikely to simultaneously produce IFNs and express RSV proteins. The modes of interference of various viruses display common, recurrent motifs, rendering the constructed model general enough to study, after readjustment of transition rates, kinetics of infection with another virus.

## Results

### Spatial stochastic model of viral infection

The constructed agent-based, spatial, stochastic model of viral infection is outlined in Fig 1. Below, we describe modeled cell components and biological processes, including virus–host cell interactions. Model derivation and parameter calibration are described in Methods. Experimental results used to parametrize the rates of processes included in the model are shown in Appendix Fig S1–S5. Model processes and rate parameters are given in Appendix Table S1.

**Figure 1.**
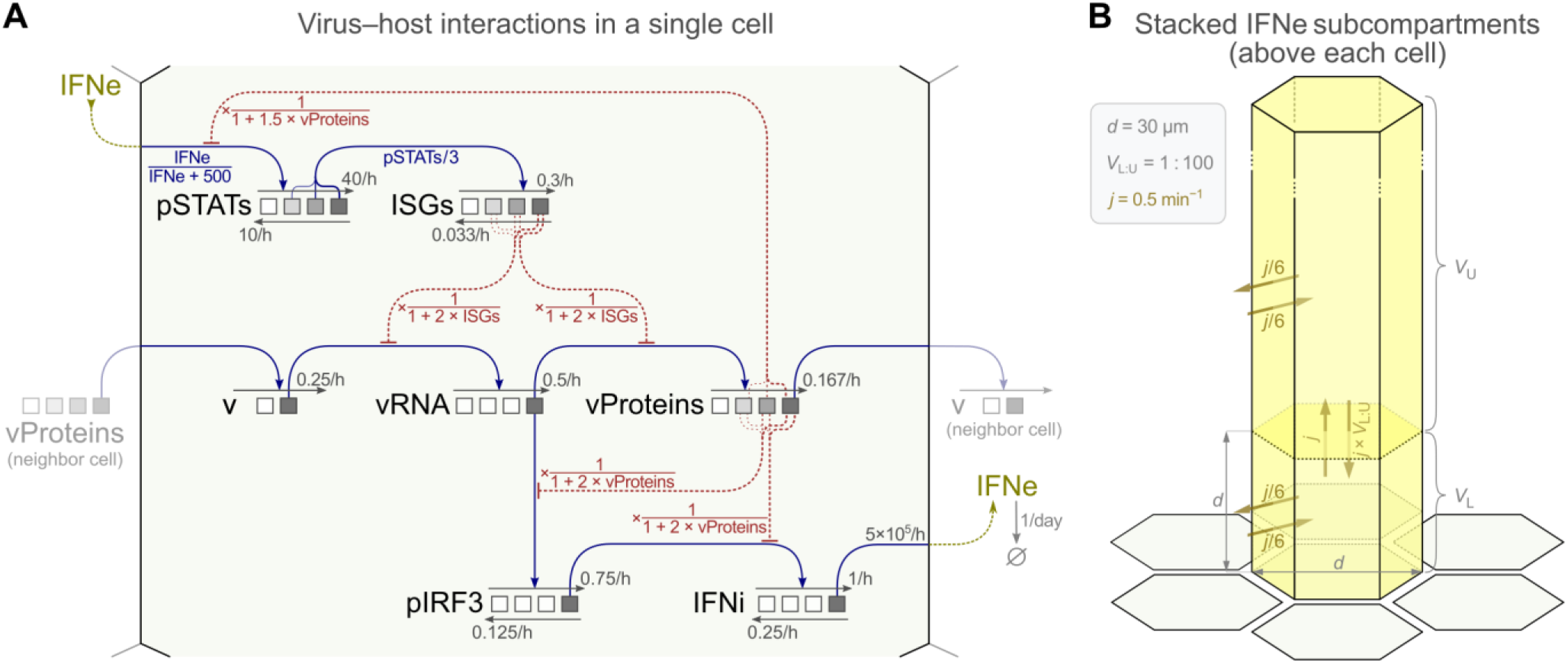
Computational model. **A** Host cell–virus interactions and adjusted kinetic rate parameter values. Black arrows indicate state transitions of seven variables characterizing the cell state: `v`, `vRNA`, `vProteins`, `pIRF3`, `IFNi`, `pSTATs`, `ISGs`. All forward transitions (thin straight arrows pointing right) are induced by other variables (as indicated by blue arrows), while the reverse transitions (thin straight arrows pointing left) have constant rates. The corresponding rate coefficients are given in gray. For forward reactions to occur, the inducing agent variable has to be maximally active, with the exception of activation of `pSTATs` (Michaelis–Menten kinetics) and activation of `ISGs` (by partially active `pSTATs`); see model reactions in Appendix Table S1. Red dashed hammer-headed arrows are inhibitory interactions: three of them emanate from `vProteins`, two emanate from `ISGs`. The expressions in red indicate how `vProteins` or `ISGs` modify the (forward) transition propensities. When an arrow has three roots (all inhibitions and `ISGs` activation), the strength of the interaction depends on the stage of activation of the inducing variable (additionally indicated by hue of the inducing variable). The extracellular interferon `IFNe` is produced at the given rate (molecules / hour) when the state of the intracellular interferon is `IFNi = 3`. `IFNe` diffuses (see panel B) and, when present in the lower subcompartment above a given cell, regulates the `pSTATs` transition rate with Michaelis–Menten kinetics. An infected cell with `vProteins = 3` infects adjacent (non-infected) cells (`v = 0` → `v = 1`) at the given rate (the infecting virions are implicit). **B** Processes related to the transport of extracellular IFN (`IFNe`) and related parameter values. `IFNe` diffuses between the lower and the upper subcompartment above each cell and between respective subcompartments above neighboring cells.

Agents (cells) are arranged in a regular two-dimensional lattice. The state of each agent-cell is fully characterized by seven discrete variables: the binary variable `v`, which indicates the cell’s infection status, and six other variables: `vRNA`, `vProteins`, `pIRF3`, `IFNi`, `pSTATs`, and `ISGs`, each assuming a discrete value from the set {`0`, `1`, `2`, `3`} and in this way describing the status of the respective intracellular biochemical species (Fig 1A). The cell state evolves as a time-continuous Markov process influenced by neighboring cells, which may infect the cell (`v = 0` → `v = 1`), and by extracellular interferon, `IFNe`.

#### Intracellular state transitions

As depicted in Fig 1A, viral infection (`v = 1`) leads to synthesis of viral RNA (increase of `vRNA`), followed by synthesis of viral proteins (increase of `vProteins`) and production of infectious virions (implicit in the model). Advancements of viral entities (`v`, `vRNA`, `vProteins`) are irreversible. The emergence of viral RNA leads to phosphorylation of IRF3 (increase of `pIRF3` at `vRNA = 3`), which in turn leads to synthesis of (intracellular) interferon (increase of `IFNi` at `pIRF3 = 3`) and its secretion. Secreted interferon is represented as a continuous variable (`IFNe`) and diffuses between subcompartments associated with individual cells (see Fig 1B and Methods for details). Extracellular interferon induces phosphorylation of STAT1 and STAT2 (represented by an increase of `pSTATs`, with saturable kinetics) that jointly trigger synthesis of proteins coded by interferon-stimulated genes (increase of `ISGs`). In the model we consider only one generic type of interferon that accounts for both type I and type III interferons (IFNβ and IFNλs, respectively; our experiments indicate that in the case of infection with RSV, IFNβ plays a decisive role, see Appendix Fig S4A and B in conjunction with Appendix Fig S1A and B, and Appendix Fig S5).

#### Virus–host cell interactions

The model includes two mutually antagonistic groups of interactions. The first group comprises a three-level inhibition of the immune response — phosphorylation of IRF3, synthesis of interferons, and phosphorylation of STAT1 and STAT2 — by viral proteins (the strengths of each inhibition increases with `vProteins`). The second group consists of two interactions, in which the accumulation of viral RNA and viral proteins is slowed down due to activity of proteins coded by interferon-stimulated genes (the strength of both inhibitions increases with `ISGs`). The presence of these two reciprocally antagonistic groups of interactions ensures the mutual inhibition of the virus and the innate immune response.

### Model simulations *vs*. immunostaining images

To characterize the incidence and co-incidence of biological entities included in the model at the single-cell level, we performed an array of experiments on A549 cell cultures infected with RSV (with or without additional IFNβ prestimulation). The cells were immunostained for RSV proteins, IRF3, phospho-Tyr701 STAT1 (p-STAT1), and IFNβ, auxiliarily stained with a DNA marker, and imaged using multi-channel confocal microscopy. In Fig 2, representative microscopy images from experiments at a multiplicity of infection (MOI) of 0.01 (Fig 2A and B) are juxtaposed with model simulations (Fig 2C). The case of an MOI of 1 is shown in Appendix Fig S6.

**Figure 2.**
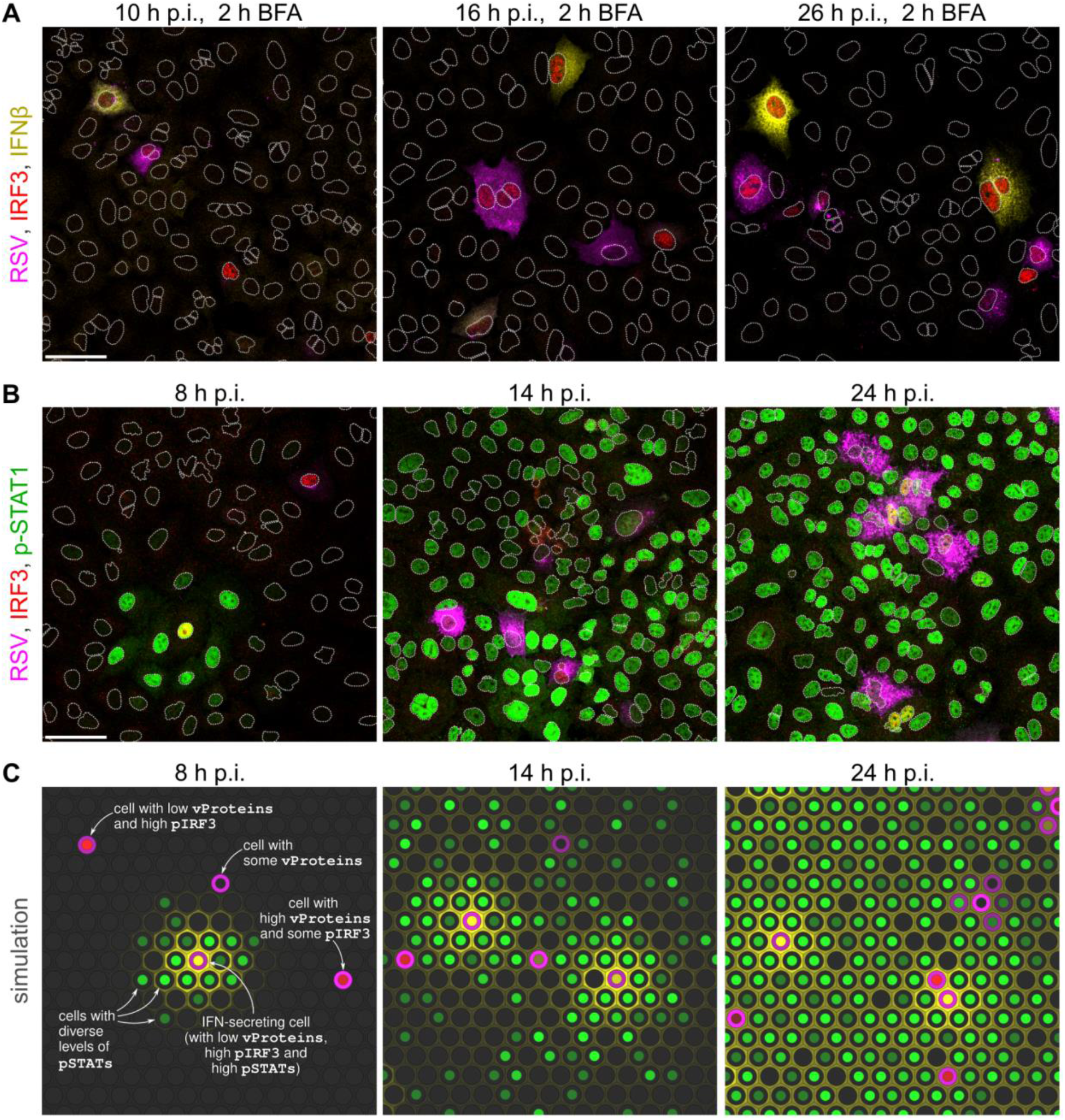
Images from experiments and snapshots from a simulation. **A, B** Overlays of images of A549 cells at three time points post infection (p.i.) with RSV at an MOI of 0.01, immunostained for: RSV proteins (polyclonal antibody) – magenta, IRF3 – red, (intracellular) IFNβ – yellow (only in panel A), phospho-Tyr701 STAT1 (p-STAT1) – green (only in panel B). The 2 h-long treatment with brefeldin A (BFA, in panel A) prevented secretion of IFNβ produced during the treatment. When phosphorylated, IRF3 translocates to the nucleus giving a discernible signal. White dotted lines are nuclear outlines determined based on DAPI counterstaining (channel not shown). Scale bars, 50 µm. **C** Snapshots from a simulation of infection at an MOI of 0.01 in a compact monolayer of cells (subpanels show fragments of a simulated 100×100 lattice). The color key corresponds to pseudocolors of immunostained proteins used in panels A, B (concentration of IFNβ in the lower subcompartment is indicated at hexagon borders; yellow color in nuclei results from mixing of red and green).

Figure 2A and B show that at 8–10 h post infection (p.i.) with RSV, the infection may manifest through IRF3 phosphorylation (p-IRF3 is translocated to the nucleus, giving a discernible nuclear IRF3 signal), expression of RSV proteins, and/or production of IFNβ (2 h-long treatment with brefeldin A prior to fixation enabled capturing IFNβ in the cytoplasm in Fig 2A). At the considered time, characteristic rings of p-STAT1 positive cells are formed, often revealing an infected cell being a local source of IFNβ (Fig 2B). In subsequent time points, 14–16 h p.i. and 24–26 h p.i., secondary infections yield small clusters of RSV proteins-producing cells. In the considered time span, IFNβ reaches all the cells and the spatial profile of p-STAT1, still concentrated around local sources of IFNβ at 14–16 h p.i., is largely homogeneous across bystander cells at 24–26 h p.i. After a high-MOI infection (Appendix Fig S6), nearly all bystander cells display p-STAT1 already at 10 h p.i., but also very likely have a virus-infected neighbor. These cells are expected to be granted less time to upregulate their ISGs prior to infection compared to the bystander cells after low-MOI infection.

As demonstrated using model trajectories (Appendix Fig S7), during a low-MOI infection, the primary infected cells exhibit fast growth of the average level of `vProteins` and have low levels of `ISGs` (Appendix Fig S7A), however the cells infected between 16 and 24 h p.i. (secondary infections) have an on average higher level of `ISGs`, which noticeably hamper production of `vProteins` (Appendix Fig S7B).

### Antagonism between RSV and the single-cell immune response RSV proteins terminate STAT activity

Immunostaining images obtained at 24 h p.i. show markedly lower levels of p-STAT1 in cells expressing RSV proteins (Fig 3A). To quantify this effect, we compared the distributions of the nuclear p-STAT1 signal in cells stratified into either expressing or not expressing RSV proteins (‘same cells’ statistics in Fig 3B). Next, we selected cells that do not express RSV proteins, divided them into either having or not having an RSV proteins-expressing cell in direct neighborhood and compared nuclear p-STAT1 between these two groups (‘neighboring cells’ statistics in Fig 3B). To characterize quantitatively the extent to which the considered distributions are disjoint, we calculated the signed Kolmogorov–Smirnov (sKS) statistic (see Methods for details). For all studied MOIs (0.01, 0.1 and 1), the value of sKS is about 0.5 for ‘same cells’ and close to 0 for the ‘neighboring cells’, which agrees well with model predictions (Fig 3C). Using the model, in Fig 3D we show that the sKS statistic for ‘same cells’ increases in time from 0 to 0.5 (stabilizing at 24 h p.i.), which indicates more pronounced virus-induced `pSTATs` deactivation at later times. The model also predicts that the value of the ‘same cells’ sKS increases with the strength of the `vProteins` ⊣`pSTATs` inhibition (Fig 3D and E), but is insensitive to variations in the strengths of the inhibition of `pIRF3` and the inhibition of `IFNi` by `vProteins` (Appendix Fig S8A). The ‘neighboring cells’ sKS statistics remain close to zero regardless of the strengths of all inhibitions originating from `vProteins` (Appendix Fig S8A and B).

**Figure 3.**
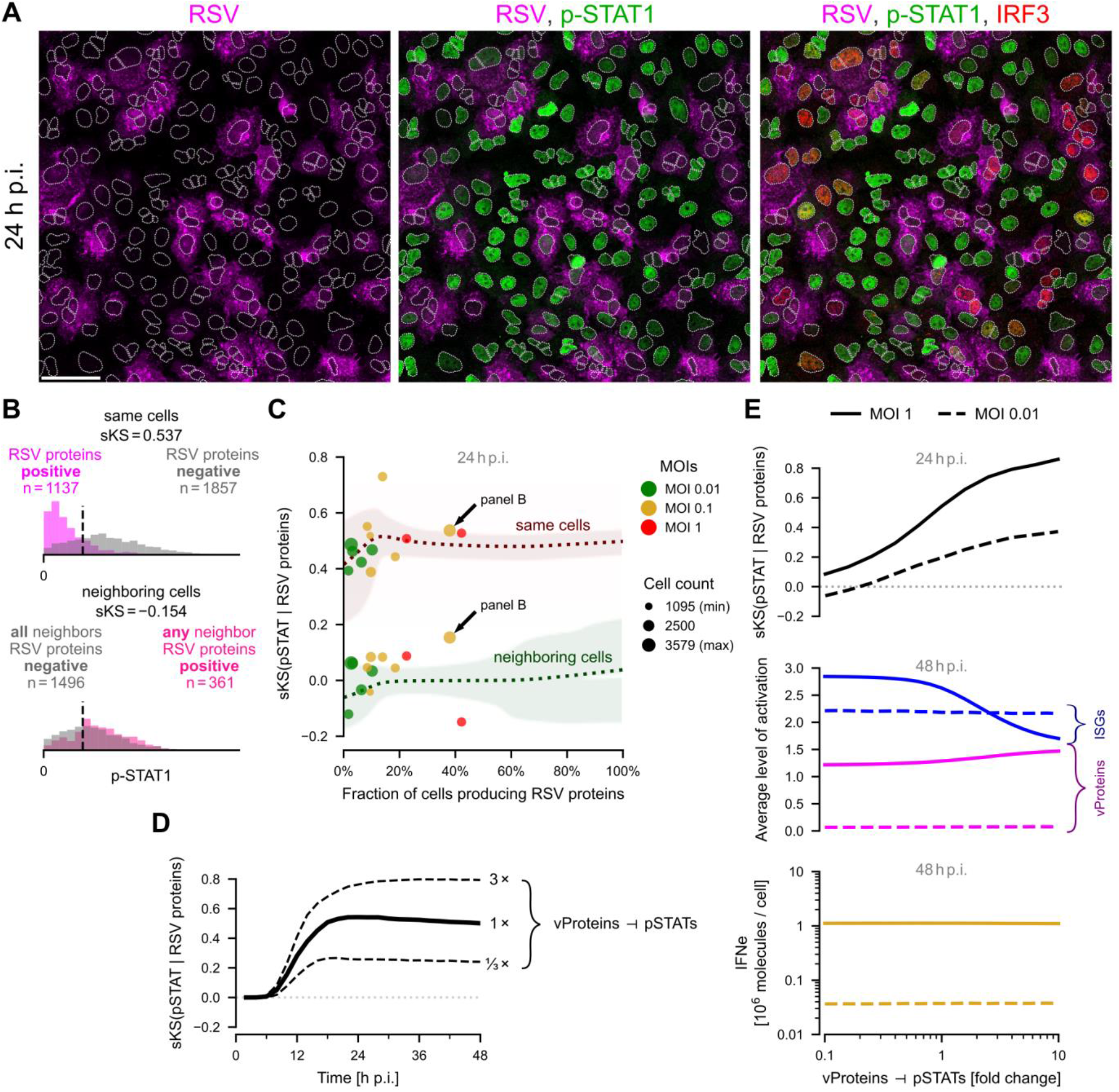
Relation between RSV proteins and STAT signaling. **A** A549 cells 24 h post infection (p.i.) with RSV at an MOI of 0.1, immunostained for RSV proteins – magenta, p-STAT1 – green, and IRF3 – red; left to right, incremental overlays. White dotted lines are nuclear outlines determined based on DAPI counterstaining (channel not shown). Scale bar, 50 µm. **B** Distributions of the mean nuclear p-STAT1 intensities in cells either expressing or not expressing RSV proteins (top subpanel, ‘same cells’) and in cells not expressing RSV proteins but either having or not a neighboring cell expressing RSV proteins (bottom subpanel, ‘neighboring cells’). Pairs of histograms, cell counts, and signed Kolmogorov–Smirnov (sKS) statistics are given for a representative experiment of an MOI of 0.1 (marked in panel C with an arrow). **C** Dependence of the ‘same cells’ sKS(pSTATs | RSV proteins) statistics and ‘neighboring cells’ sKS(pSTATs | RSV proteins) statistics on the fraction of RSV proteins-expressing cells at 24 h p.i. Marker sizes correspond to the number of cells quantified in each experiment. Experimental results are juxtaposed with model predictions (dotted lines enveloped within 95% CrI computed for *n* = 2500 cells). **D** ‘Same cells’ sKS(pSTATs | RSV proteins) over the period of two days post infection at an MOI of 0.1, according to the model. The dashed lines were obtained at different strengths of the `vProteins` ⊣`pSTATs` inhibition. **E** Model-based analysis of the influence of the `vProteins` ⊣`pSTATs` inhibition: ‘Same cells’ sKS(pSTATs | RSV proteins) (upper subpanel, at 24 h p.i.), average `ISGs` and `vProteins` states (middle subpanel, at 48 h p.i.), average level of extracellular interferon `IFNe` (lower subpanel, at 48 h p.i.).

Surprisingly, the model indicates that the strength of `pSTATs` inhibition by `vProteins` − although influencing the status of `pSTATs` in `vProteins`-expressing cells − very weakly influences the overall spread of infection (Fig 3E). For an MOI of 1 (but not for an MOI of 0.01) and stronger `vProteins` ⊣`pSTATs` inhibition, the average status of `ISGs` is lowered, but this leads to a very modest increase of the average status of `vProteins` at 48 h p.i, and does not influence extracellular IFN accumulation. This is because, in individual cells, inhibition of `pSTATs` and consecutive reduction of the build-up of `ISGs` typically occurs after the (irreversible) accumulation of `vProteins`, and is thus not advantageous to the virus.

### RSV proteins inhibit IRF3 activation

Viral RNA is required for synthesizing RSV proteins as well as for activating IRF3; these two processes, however, may but do not need to occur simultaneously in the same cell. Accordingly, in our immunostaining images there are cells expressing RSV proteins with and without active IRF3, as well as cells with active IRF3 but devoid of RSV proteins (Fig 4A). Imaging data collected at different time points (16, 20, 24, 36, 40, and 48 h p.i.) and various initial MOIs (0.01, 0.1, and 1) show that approximately 40% of RSV proteins-expressing cells have active IRF3, with a good agreement with the model (Fig 4B). Conversely, approximately 50–80% of cells with active IRF3 express RSV proteins (Fig 4C). In experiments the percentage of IRF3-active cells is somewhat larger for a larger fraction of cells expressing RSV proteins, but this finding is not reproduced by our model. Additionally, in Fig 4B and C, we observe high variability between experimental replicates, indicating that the counterbalance between the strength of the innate immune response (indicated by the fraction cells with active IRF3) and virus spread (indicated by the fraction of cells with RSV proteins) is heterogeneous and sensitive to experimental conditions.

**Figure 4.**
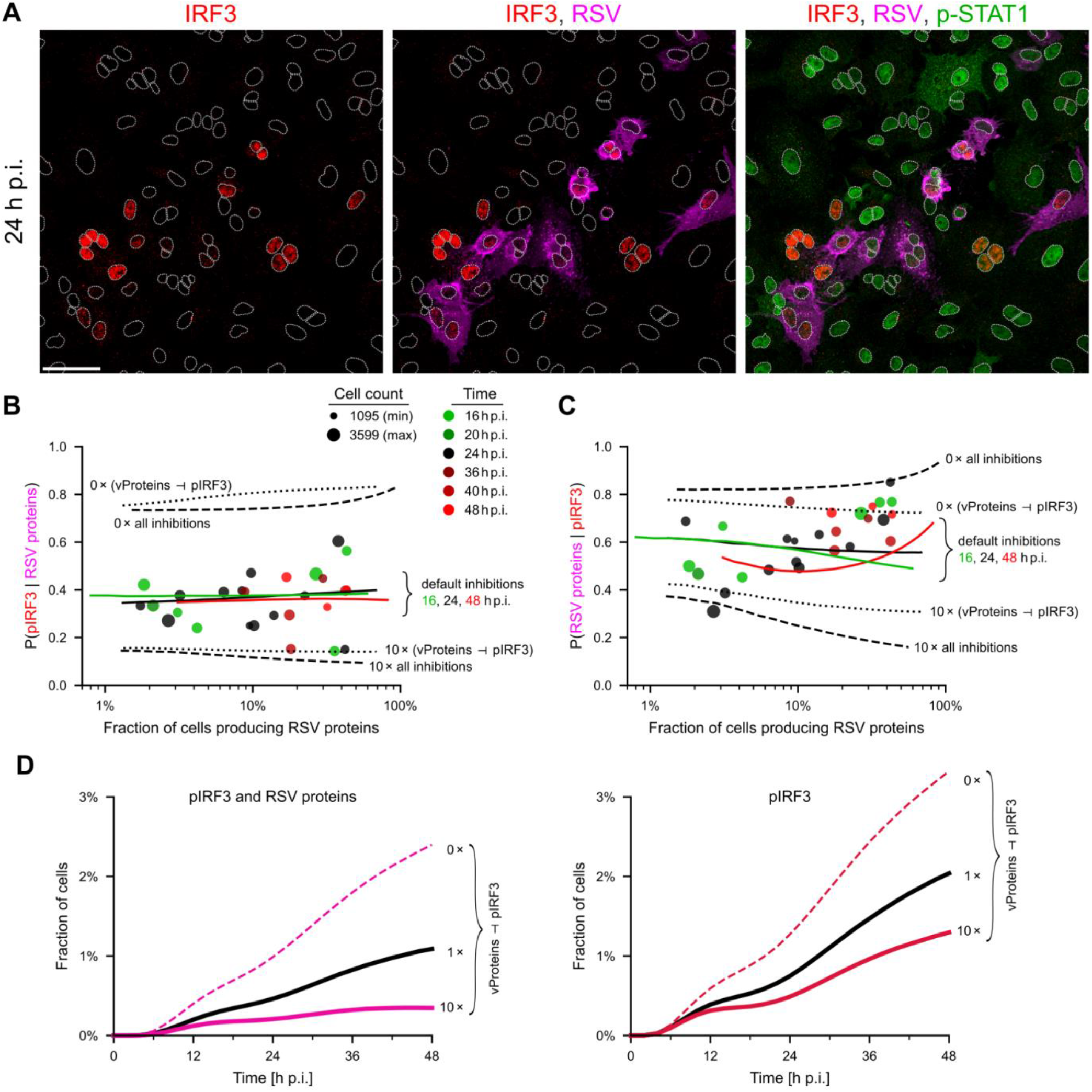
Relation between expression of viral proteins and IRF3 activity. **A** Incremental overlays of images of A549 cells 24 h post infection (p.i.) with RSV at an MOI of 0.1, immunostained for IRF3 – red, RSV – magenta, and p-STAT1 – green. White dotted lines are nuclear outlines determined based on DAPI counterstaining (channel not shown). Scale bar, 50 µm. **B** Conditional probability *P*(pIRF3 | RSV proteins) as a function of the percentage of RSV proteins-expressing cells. Experiments (at 16, 20, 24, 36, 40, 48 h p.i.) – disks, model (at 16, 24, 48 h p.i.) – lines (line points were computed for a specific MOI, resulting in a given proportion of RSV-positive cells; results are averages over 1000 stochastic simulations on the 100×100 lattice). Dashed lines show model predictions (at 24 h p.i.) without all 5 inhibitions or with 10 times stronger inhibitions (as shown in the figure). Dotted lines show model predictions (at 24 h p.i.) without the `vProteins` ⊣`IRF3` inhibition or with 10 times stronger inhibition. **C** Conditional probability *P*(RSV proteins | pIRF3), notation as in panel B. **D** Proportion of the (IRF3 & vProtein)-positive cells (left) and IRF3-positive cells (right) as a function of time for a default strength of the `vProteins` ⊣ `pIRF3` inhibition, no inhibition, and the inhibition 10 times stronger than default.

To show the importance of inhibitions, we include model predictions with the `vProteins` ⊣ `pIRF3` inhibition and with all inhibitions either turned off or made 10-fold stronger (Fig 4B and C). This analysis highlights the key role of the `vProteins` ⊣ `pIRF3` inhibition in regulation of the proportions of cells co-expressing active IRF3 and RSV proteins with respect to cells expressing RSV proteins (Fig 4B) or active IRF3 (Fig 4C). According to the model, the fraction of cells that display both RSV proteins and IRF3 activity increases in time (as infection progresses) and substantially decreases with the strength of inhibition of `pIRF3` by `vProteins` (Fig 4D, left subpanel). Consequently the overall fraction of IRF3-active cells is lower for stronger inhibition (Fig 4D, right subpanel).

### Mutual exclusion between expression of RSV proteins and synthesis of IFNβ

Viral RNA triggers production of IFN (via activation of IRF3) and is required for synthesis of viral proteins. However, as we can see in immunostaining images in Fig 5A, accumulation of IFN and RSV proteins rarely coincides in the same cell. The analysis of immunostaining images indicates that about 10% of cells expressing RSV proteins exhibit IFN accumulated during 2 hours of brefeldin A treatment that blocks IFN secretion (Fig 5B). In turn, 30–40% of the cells exhibiting accumulated IFN have RSV proteins (Fig 5C). Both percentages do not change significantly with initial MOIs (0.01, 0.1, and 1) and over time (experimental time points: 16, 20, 24, 36, 40, and 48 h p.i.), and are in good agreement with the model. As previously in Fig 4B and C, in Fig 5B and C we observe high variability between experimental replicates.

**Figure 5.**
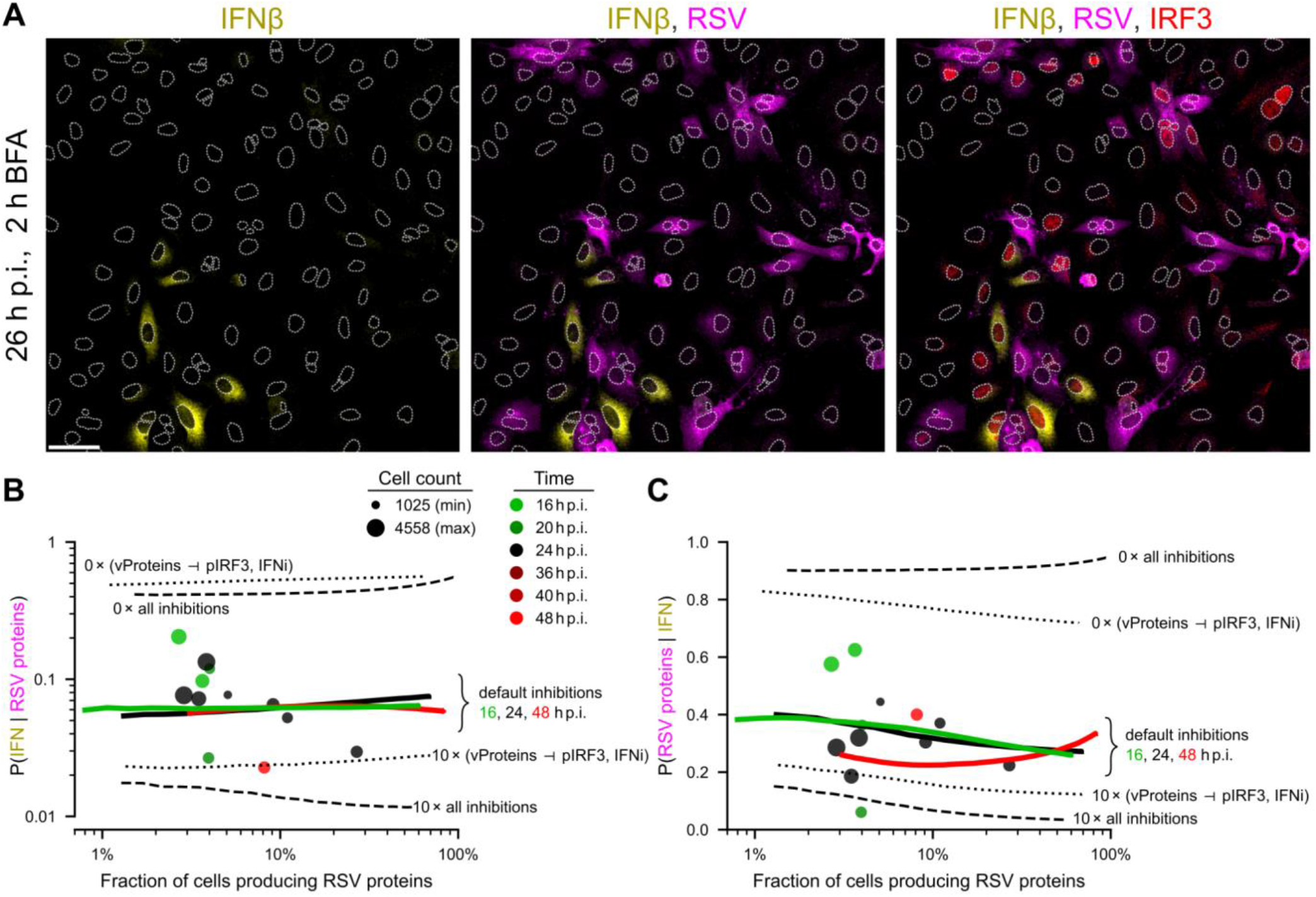
Mutually exclusive expression of RSV proteins and synthesis of IFNβ. **A** Incremental overlays of images of A549 cells 26 hours post infection (h p.i.) with RSV at MOI 0.1, immunostained for IFNβ – yellow, RSV – magenta, and IRF3 – red. The 2 hour-long treatment with brefeldin A (BFA) prevents secretion of IFNβ produced by some cells during the treatment. White dotted lines are nuclear outlines determined based on DAPI counterstaining (channel not shown). Scale bar, 50 µm. **B** Conditional probability *P*(IFN | RSV proteins) as a function of the percentage of RSV proteins-expressing cells. Experiments (at 16, 20, 24, 36, 40, 48 h p.i.) – disks, model (at 16, 24, 48 h p.i.) – lines (line points were computed for a specific MOI, resulting in a given proportion of RSV-positive cells; results are averages over 1000 stochastic simulations on the 100×100 lattice). Dashed lines show model predictions (at 24 h p.i.) without all 5 inhibitions or with 10 times stronger inhibitions (as shown in the figure). Dotted lines show model predictions (at 24 h p.i.) without `vProteins` ⊣ `pIRF3` and `vProteins` ⊣ `IFNi` inhibitions, or with these two inhibitions 10 times stronger. **C** Conditional probability *P*(RSV proteins | IFN), notation as in panel B.

The model predictions with inhibitions `vProteins` ⊣ `pIRF3` and `vProteins` ⊣ `IFNi`, and with all inhibitions either turned off or made 10-fold stronger (Fig 5B and C), show the key role of the `vProteins` ⊣ `pIRF3` and `vProteins` ⊣ `IFNi` inhibitions in achieving correct proportions of cells co-expressing IFN and RSV proteins with respect to cells expressing RSV proteins (Fig 5B) or IFN (Fig 5C).

### Model-based analysis of the influence of inhibitory interactions on the progression of viral infection

The effect of antagonistic inhibitions from `vProteins` to `pIRF3` and `IFNi`, and from `ISGs` to `vRNA` and `vProteins` (Fig 6A) on the progression of the infection is further studied using the validated model. We focus on the case of MOI = 0.01, in which the immune response is granted time to develop.

**Figure 6.**
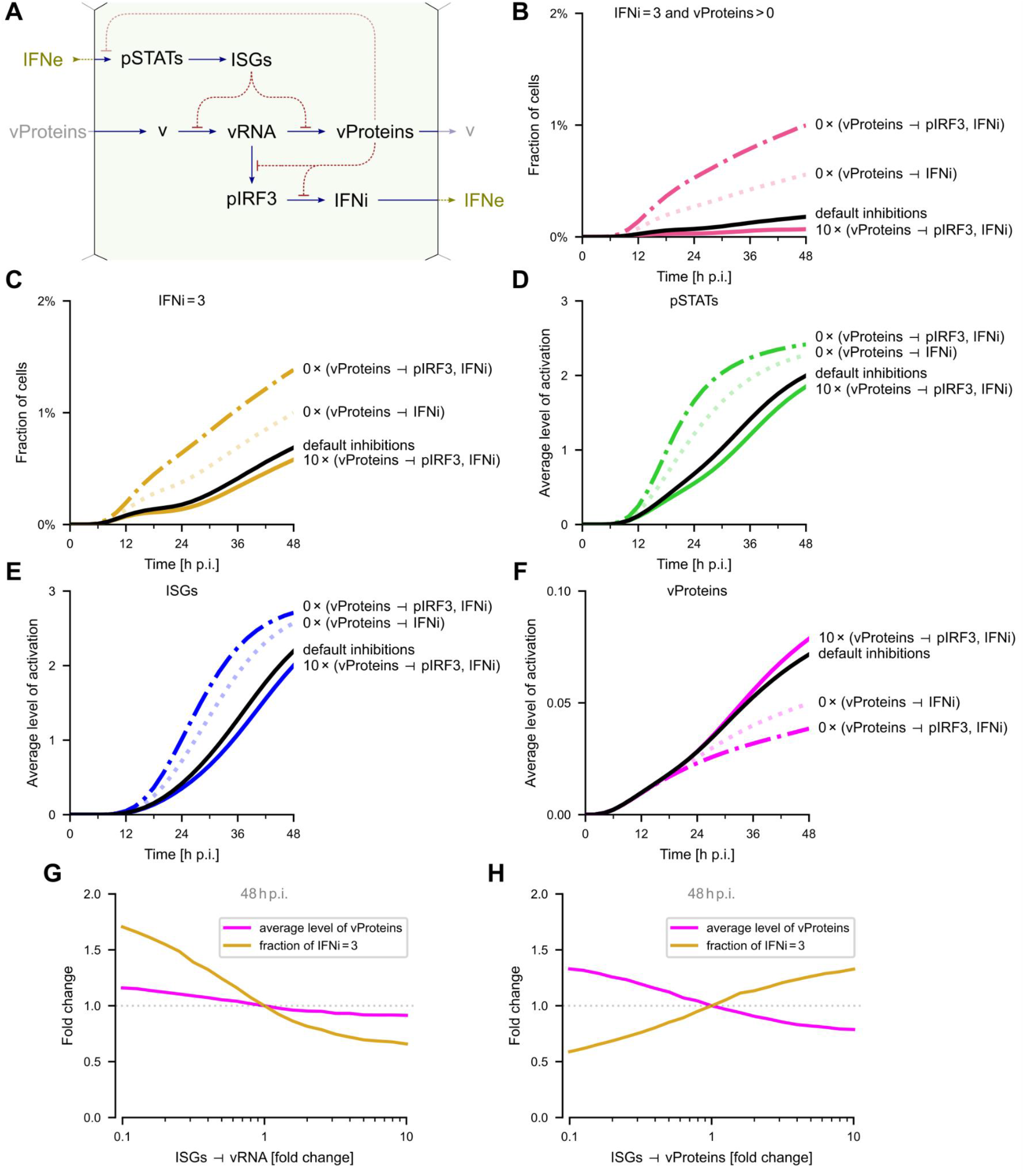
Influence of inhibitions on the progression of viral infection (model). **A** Simplified scheme of antagonistic interaction present in the model. **B–E** (**B**) Fraction of cells producing `IFNe` (`IFNi = 3`) and `vProteins` (`vProteins > 0`) simultaneously, (**C**) fraction of cells producing `IFNe`, (**D**) average activation of `pSTATs`, (**E**) average activation of `ISGs`, (**F**) fraction of cells producing `vProteins`; all as a function of time for an MOI of 0.01 for default inhibition strengths (black line), inhibitions `vProteins` ⊣ `IRF3` and `vProteins` ⊣ `IFNi` 10 times stronger (thick line), and turned off (dashed, dotted and dash-dotted lines). **G** Change of the average level of `vProteins` and of the fraction of cells producing `IFNe` as a function of the `ISGs` ⊣ `vProteins` inhibition, at 48 h p.i. at an MOI of 0.01. **H** Change of the average level of `vProteins` and of the fraction of cells producing `IFNe` as a function of the `ISGs` ⊣ `vRNA` inhibition, at 48 h p.i. at an MOI of 0.01.

As shown in Fig 6B, the fraction of cells producing jointly IFN and RSV proteins grows in time as the infection spreads, but remains below 0.2% even at 48 h p.i. This fraction is controlled by the strengths of inhibitions `vProteins` ⊣ `pIRF3` and `vProteins` ⊣ `IFNi`. When these inhibitory interactions are 10 times stronger, the fraction is close to 0, whereas their removal increases the fraction fivefold, to 1%. Correspondingly, the removal of these inhibitory interactions increases the overall fraction of cells producing IFN (at 48 h p.i.) more than twofold, to above 1% (Fig 6C). However, when 10-fold stronger inhibitory interactions are assumed, the relative effect on the fraction of IFN-producing cells is negligible, since only about 10% of cells producing IFN have `vProteins` (consistently with data in Fig 5B). The effect of removing inhibitory interactions `vProteins` ⊣ `pIRF3` and `vProteins` ⊣ `IFNi` is forwarded through subsequent tiers of the immune response leading to an increase of the levels of `pSTATs` (Fig 6D) and `ISGs` (Fig 6E). The most significant increase is observed, respectively, at about 20 and 30 h p.i. Consequently, induction of a more pronounced antiviral state at about 30 hours (epitomized by an increased level of `ISGs`) leads to a decrease of `vProteins`-producing cells after that time point (Fig 6F). In summary, the analysis presented in Fig 6B–F dissects regulatory steps in which the virus enhances its propagation by suppressing the innate immune response.

Next, we analyze how the strengths of inhibitory interactions mediated by the immune response, `ISGs` ⊣ `vProteins` and `ISGs` ⊣ `vRNA`, influence the propagation of infection and activation of the immune response. As shown in Fig 6G, an increase of the strength of the `ISGs` ⊣ `vProteins` inhibition leads to a decrease of the fraction of `vProteins`-producing cells and to a simultaneous increase of the number of interferon-producing cells. Clearly, this inhibitory interaction attenuates viral infection and promotes interferon signaling. The effect of inhibitory interaction `ISGs` ⊣ `vRNA` is different; an increase of its strength leads to a moderate decrease of the fraction of `vProteins`-producing cells, but also to a substantial decrease of interferon-producing cells (Fig 6H). This is because viral RNA is necessary for both the synthesis of viral proteins and activation of the IRF3 → IFN pathway.

### Influence of DVGs on virus spread

In the model we have not accounted for potential formation of defective viral particles with incomplete genomes that are capable of activating IRF3 (triggering IFN synthesis), but neither produce infectious virions nor nonstructural proteins that would inhibit IFN synthesis. In experimental conditions the presence of DVGs depends on viral amplification (Sun *et al*, 2015) and although we cannot rule out their presence in our experiments, the obtained results can be explained by the model that neglects their presence. Since DVGs are known to be important drivers of the immune response (Vignuzzi & López, 2019), we analyze theoretically the consequence of their presence on IFN synthesis and virus spread.

For the sake of simplicity we assume that there is a fraction of cells infected solely with DVGs (*f*_DVG_) which are capable of activating IRF3 but, in the absence of replication-competent helper virions, cannot express viral proteins. In Fig 7A we show a relatively modest increase of `IFNe` and about twofold decrease of the average `vProteins` level as *f*_DVG_ increases from 0 to 30%. This surprisingly weak influence is the consequence of the fact that for the default model parameters, cells are able to express IFN before viral proteins are produced. However, when the `vProteins` forward rate is increased threefold, the effect of DVGs infected cells is much stronger, leading to fourfold reduction of the level of `vProteins`. Overall, this analysis indicates that DVGs can be critical drivers of innate responses, but only in the case of viruses that are capable of quickly terminating synthesis of IFN.

**Figure 7.**
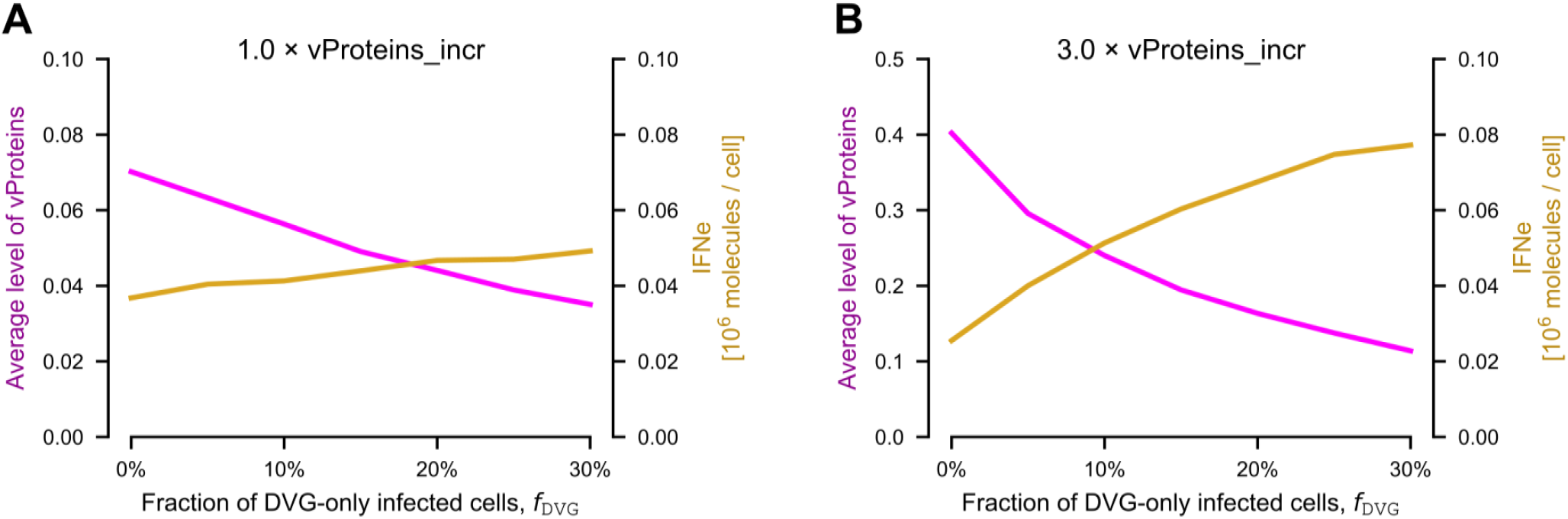
Influence of DVGs on IFN expression and viral progeny (model). **A** The average status of `vProteins` and `IFNe` as a function of the percentage of cells infected solely by DVG at 48 h p.i. Initial infection MOI: 0.01. **B** Same as in panel A, but with the `vProteins` forward rate coefficient (`vProteins_inc`) increased 3-fold. Note the different range on the left vertical axis.

## Discussion

Viral infection, when viewed at the cell-population level, leads to both multiplication of viral particles and triggering of the innate immune response. The intuitively expected antagonism between these two processes can be investigated at the single-cell level. Based on cell-population and single-cell experiments involving RSV, we constructed and constrained a stochastic, agent-based model that elucidates this mutual antagonism. The model allowed us to investigate correlations between the levels of key proteins in the same cell as well as in adjacent cells, which enabled quantitative comparison with spatial patterns observed in immunostaining images.

Both the experimental data and the model indicate that while viral RNA triggers the innate immune response, viral proteins inhibit this response at 3 levels: by blocking activation of IRF3, by blocking synthesis of IFN, and by inhibiting STATs activation. The first two interactions promote the spread of infection, whereas, as suggested by the model, the last one has a minimal influence on virus spread. The lack of influence of the STAT signaling inhibition by RSV (nonstructural) proteins on RSV replication observed in the model suggests that this interaction may be implicated in regulatory processes not included in the model.

The innate immune response inhibits viral propagation through paracrine IFN signaling, which activates STAT signaling which, in turn, upregulates expression of ISGs. Once a cell becomes infected, ISGs slow down synthesis of RSV proteins, giving more time for activation of IRF3 and synthesis of IFN. However, because the process of RSV protein synthesis is irreversible, at some point, RSV proteins accumulate and terminate both the activity of IRF3 and synthesis of IFN. A consequence of the `vProteins` ⊣ `pRF3` and `vProteins` ⊣ `IFN` inhibitions is that only about 40% of viral proteins-expressing cells have active IRF3 and as little as 10% of such cells produce IFN.

This picture is changed in the presence of DVGs; the model indicates that the fraction of cells infected solely by DVGs can, due to the lack of viral protein expression, serve as a source of IFN and in that way limit virus spread. The effect is modest for default model parameters but significant when faster accumulation of viral proteins is assumed—this implies that DVGs can be critical drivers of interferon-mediated responses, but only in the case of viruses that otherwise would effectively inhibit interferon synthesis.

It is known that RSV (using NS1 and NS2) effectively blocks cell apoptosis (Bitko *et al*, 2007), which is supported by our immunostaining images taken at 48 h p.i., indicating that even in the foci of infected regions, cells remain confluent. For many other viruses, however, apoptosis is known to be an important factor limiting the spread of infection (Barber, 2001). Since our model is calibrated for RSV, it does not account for apoptosis. As a consequence, interferon signaling (by triggering antiviral state in not yet infected cells) can only slow down virus spread, but cannot suppress it. Induction of apoptosis can change this picture through elimination of infected cells before they produce new virions.

In summary, single-cell experimental data and the proposed stochastic, agent-based model demonstrate the importance of reciprocally inhibitory interactions between the virus and the innate immune response. Because of the irreversible progression of virus replication (not limited by cell apoptosis), these interactions do not lead to a typical bistability; instead, they result in divergent single-cell trajectories. A single infected cell may either rapidly initiate a cascade of new infections or become a transient source of IFN, allowing neighboring cells to enter an antiviral state and slow down the virus spread. As a consequence, when the respiratory epithelium is infected by a small number of virions, heterogeneity of single-cell responses introduced by noise and mutually inhibiting interactions may lead to different outcomes of infection at the organismal level.

## Methods

### Experimental methods

#### Cell lines and cell culture

The A549, HeLa (used for RSV proliferation), and HEK-293T (used for production of lentiviral particles) cell lines were purchased from ATCC. A549 cells were cultured in F-12K basal medium. HeLa and HEK-293T cells were maintained in Dulbecco’s modified Eagle’s medium (DMEM). Both media were supplemented with 10% FBS and a penicillin–streptomycin antibiotic solution. Cells were cultured under standard conditions (37 °C with 5% CO_2_) in a series 8000 WJ incubator (Thermo Fisher Scientific) and kept in monolayers up to 90% confluency.

For STAT1 and STAT2 gene knockouts, a CRISPR lentiviral vector system was used. Commercially available sgRNAs sets were purchased from Applied Biological Materials (catalog numbers, K0002501 and K2300601, respectively). STAT1 knockout was obtained by targeting sequence TGATCCAAGCAAGCATTGGG in exon 9 of STAT1 gene; STAT2 knockout was obtained by targeting sequence AGCTCCCATTGACCACGGGT in exon 8 of STAT2 gene. Two days after transfection of HEK-293T cells with plasmids encoding Cas9 nuclease (catalog number K002; Abm), lentivirus packaging particles (catalog number LV053; Abm), and mentioned earlier of each single sgRNA, the media with lentiviral particles were collected, enriched with 8 mg/mL Polybrene, and filtered through 0.45-mm syringe filters. Next, each lentiviral supernatant was used to transduce A549 wild-type cells, which were subcultured at low confluence (seeding density of 5×10^4^ cells per 30-mm dish). After another 2 days, A549 cells were subjected to selection with 800 mg/mL of G418 for 10 days and seeded into a 96-well plate to obtain single-cell colonies. Clones were validated by Western blotting of p-STAT1 and p-STAT2 in response to IFNβ. Additionally, the KO clones selected for further experiments were verified by sequencing.

#### RSV amplification and isolation

Respiratory syncytial virus strain A2 was purchased from ATCC and amplified in HeLa cells. The cells were seeded into 225-cm^2^ tissue culture flasks (Falcon) and cultured as described above for 2 to 3 days until they reached 90% confluence. The virus growth medium (DMEM with 2% FBS) and a virus dilution (with a target MOI of ∼0.01) were prepared on the day of infection. The culture medium was removed and cells were washed once with phosphate-buffered saline (PBS) and overlaid with 10 mL of the inoculum. Virus was allowed to adsorb to cells for 2 h at 37 °C, with occasional stirring. Next, additional virus growth medium was added to a total volume of 40 mL per flask. Infected cells were cultured for 3– 6 days at 37 °C, until the development of cytopathic effects was observed in at least 80% of the cells. Virus-containing culture fluid was then collected and clarified by centrifugation at 3,000 *g* at 4 °C for 20 min. Next, virus particles were precipitated by adding 50% (wt/vol) polyethylene glycol 6000 (PEG 6000) (Sigma-Aldrich) in NT buffer (150 mM NaCl, 50 mM Tris-HCl [pH 7.5]) to a final concentration of 10% and stirring the mixture gently for 90 min at 4 °C. Virus was centrifuged at 3,250 *g* for 20 min at 4 °C and re-centrifuged after the removal of the supernatant to remove the remaining fluid. The pellet was suspended in 1 mL 20% sucrose in the NT buffer, aliquoted and stored at −80 °C.

#### RSV quantification

The virus concentration was quantified using an immunofluorescence protocol. HeLa cells were seeded onto microscopic coverslips and cultured upon reaching 90 to 100% confluence. Serial dilutions of virus samples were prepared in the virus growth medium in a range of 10^−3^ to 10^−7^. After washing with PBS, cells were overlaid with diluted virus (for each virus dilution, two coverslips were used). Virus was allowed to adhere for 2 h and occasionally stirred. Afterwards, the virus-containing medium was removed, and cells were overlaid with fresh virus growth medium and cultured for 24 h. Next, cells were washed with PBS and fixed with 4% formaldehyde for 20 min at room temperature. Cells were stained using a standard immunofluorescence protocol with an anti-RSV fusion glycoprotein antibody (catalog number ab43812; Abcam). Cells containing stained viral proteins were counted using a Leica TCS SP5 X confocal microscope. The virus concentration was calculated using the following formula: (average number of infected cells)/(dilution factor × volume containing virus added) = infectious particles/mL.

#### Compounds and stimulation protocols

Human IFNβ 1a was purchased from Thermo Fisher Scientific (catalog number PHC4244) and prepared according to the manufacturer’s instructions. For cell stimulation, interferons were further diluted to the desired concentration in F-12K medium supplemented with 2% FBS. The decreased FBS content was used to prevent the inhibition of viral attachment and entry at the second stage of experiments. For interferon-and-virus experiments, the cell culture media were exchanged for interferon-containing or control media at time zero and were not removed until the end of the experiment. Appropriately diluted virus was added in small volumes (less than 10 μl) directly into the wells. The even distribution of virus across the cell population was aided by the intermittent rocking of the plate for 2 h. For intracellular IFNβ visualization, a brefeldin A solution (catalog number 00-4506-51; Invitrogen) was added 2 h prior to cell fixation.

#### Western blotting

At the indicated time points, cells were washed twice with PBS, lysed in Laemmli sample buffer containing dithiothreitol (DTT), and boiled for 10 min at 95 °C. Equal amounts of each protein sample were separated on 4% to 16% Mini-Protean TGX stain-free precast gels using the Mini-Protean Tetra cell electrophoresis system (Bio-Rad). Upon the completion of electrophoresis, proteins were transferred to a nitrocellulose membrane using wet electrotransfer in the Mini-Protean apparatus (400 mA for 50 min). The membrane was rinsed with TBST (Tris-buffered saline [TBS] containing 0.1% Tween 20; catalog number P7949, Sigma-Aldrich) and blocked for 1 h with 5% BSA–TBS or 5% nonfat dry milk. Subsequently, the membranes were incubated with one of the primary antibodies diluted in a 5% BSA–TBS buffer at 4 °C overnight. After thorough washing with TBST, the membranes were incubated with secondary antibodies conjugated to a specific fluorochrome (DyLight 800; Thermo Fisher Scientific) or horseradish peroxidase (HRP-conjugated polyclonal anti-mouse/anti-rabbit immunoglobulins; Dako) diluted 1:5,000 in 5% non-fat dry milk–TBST for 1 h at room temperature. The chemiluminescence reaction was developed with the Clarity Western ECL system (Bio-Rad). For GAPDH detection, hFAB rhodamine anti-GAPDH primary antibody (Bio-Rad) was used. Specific protein bands were detected using the ChemiDoc MP imaging system (Bio-Rad).

#### Immunostaining

For staining of intracellular proteins, cells were seeded onto 12-mm round glass coverslips, which were previously washed in 60% ethanol–40% HCl, thoroughly rinsed with water, and sterilized. After stimulation, cells on the coverslips were washed with PBS and immediately fixed with 4% formaldehyde (20 min at room temperature). Cells were then washed 3 times with PBS and incubated with 100% cold methanol for 20 min at −20 °C. After washing with PBS, coverslips with cells were blocked and permeabilized for 1.5 h with 5% BSA (Sigma-Aldrich) with 0.3% Triton X-100 (Sigma-Aldrich) in PBS at room temperature. Subsequently, coverslips with cells were incubated with primary antibodies diluted in a blocking solution overnight at 4 °C. Cells were then washed 5 times with PBS and then appropriate secondary antibodies conjugated with fluorescent dyes were added in a blocking solution for 1 h at room temperature. Subsequently, cells were washed with PBS and their nuclei were stained with 200 ng/mL 4’,6-diamidino-2-phenylindole (DAPI) (Sigma-Aldrich) for 10 min. After a final wash in MilliQ water, coverslips with stained cells were mounted onto microscope slides with a Vectashield Vibrance Antifade Mounting Medium (Vector Laboratories). Multichannel fluorescence images were acquired with a Leica TCS SP5 X confocal microscope.

### Antibodies

#### Antibodies for Western blotting

##### Primary antibodies

anti-phospho-STAT1 (Tyr701) (clone 58D6) (catalog number 9167; Cell Signaling Technologies; 1:1000); anti-phospho-STAT2 (Tyr690) (clone D3P2P) (catalog number 88410; Cell Signaling Technologies; 1:1000); anti-phospho-IRF3 (Ser396) (clone 4D4G) (catalog number 4947; Cell Signaling Technologies; 1:1000); anti-IRF3 (clone D6I4C) (catalog number 11904; Cell Signaling Technologies; 1:1000); anti-RIG-I (clone D14G6) (catalog number 3743; Cell Signaling Technologies; 1:1000); anti-STAT1 (catalog number 610116; BD Biosciences; 1:1000), anti-STAT2 (catalog number PAF-ST2; R&D Systems; 1:1000); anti-PKR (clone B-10) (catalog number sc-6282; Santa Cruz Biotechnology; 1:1000); anti-RNase L (clone E-9) (catalog number sc-74405; Santa Cruz Biotechnology; 1:1000); anti-OAS1 (clone F-3) (catalog number sc-374656; Santa Cruz Biotechnology; 1:1000), anti-Respiratory Syncytial Virus (clone 2F7) (catalog number ab43812; Abcam; 1:1000); hFAB Rhodamine anti-GAPDH (catalog number 12004168; Bio-Rad; 1:10,000).

##### Secondary antibodies

goat anti-rabbit IgG (H+L), DyLight 800 4X PEG (catalog number SA5-35571; Thermo Fisher Scientific; 1:10,000); goat anti-mouse IgG (H+L), DyLight 800 4X PEG (catalog number SA5-35521; Thermo Fisher Scientific; 1:10,000); StarBright Blue 700 goat anti-mouse IgG (catalog number 12004159; Bio-Rad; 1:10,000), rabbit anti-goat immunoglobulins/HRP (catalog number P0449; Agilent; 1:10,000).

#### Antibodies for immunostaining

##### Primary antibodies

anti-phospho-STAT1 Tyr701 (clone 58D6) (catalog number 9167; Cell Signaling Technologies; 1:1000); anti-IRF-3 (clone D-3) (catalog number sc-376455; Santa Cruz Biotechnology; 1:500), anti-Respiratory Syncytial Virus (catalog number ab20745; Abcam; 1:1000), anti-human interferon beta (catalog number MAB8142; R&D Systems; 1:100).

##### Secondary antibodies

donkey anti-rabbit IgG (H+L), Alexa Fluor 488 conjugate (catalog number A-21206; Thermo Fisher Scientific; 1:1000); donkey anti-mouse IgG (H+L), Alexa Fluor 555 conjugate (catalog number A-31570; Thermo Fisher Scientific; 1:1000); donkey anti-goat IgG (H+L), Alexa Fluor 633 conjugate (catalog number A-21082; Thermo Fisher Scientific; 1:1000).

### Computational methods and modeling

#### Image quantification

Confocal images obtained from immunostaining were analyzed using our in-house software (https://pmbm.ippt.pan.pl/software/shuttletracker). Nuclear regions were detected based on DAPI staining. The nuclei that were partially out of frame or mitotic were excluded from the analysis; outlines of overlapping nuclei were split based on geometric convexity defects when possible. To characterize cells with respect to the levels of RSV proteins, nuclear IRF3 (as a proxy of p-IRF3), accumulated IFNβ, and p-STAT1, we used image features that were visually checked and confirmed to adequately capture the immunofluorescence signal in a number of fields of view in several experiments. Specifically, cells were classified as: RSV proteins positive/negative — based on both the mean intensity of pixels of a perinuclear ring (after rejecting ⅓ of the brightest and ⅓ of the darkest pixels) and the sum of pixel intensities in the nuclear region and in the sigmoidally weighted nuclear halo around the nucleus in its corresponding Voronoi tile (2 features); p-IRF3 positive/negative — based on both the mean intensity of the nuclear region (after rejecting 10% brightest and 10% darkest pixels) and nuclear dominance (calculated as the difference between the mean nuclear and the mean perinuclear ring intensity normalized by the sum of these mean intensities, all calculated after rejecting 10% brightest and 10% darkest pixels; 2 features); IFNβ positive/negative — based on both the mean intensity of the 50% brightest pixels in the perinuclear ring and the sum of pixel intensities in the nuclear region and in the sigmoidally weighted nuclear halo around the nucleus in its corresponding Voronoi tile (2 features); p-STAT1 was quantified as the mean intensity of the nuclear region (after rejecting 10% brightest and 10% darkest pixels; 1 feature, not subjected to thresholding). Thresholds used to binarize the features of RSV proteins, IRF3, and IFNβ were selected to correctly identify cells visually identified as RSV proteins, p-IRF3, and IFNβ positive/negative in a number of fields of view in several experiments.

To express the extent to which the two distributions are disjoint, we calculated the signed Kolmogorov–Smirnov (sKS) statistic. The absolute value of sKS is that of the standard Kolmogorov– Smirnov statistics, whereas the sign additionally informs about the relative locations of two (empirical) probability densities (shown in histograms as probability density functions, PDFs). We used the notation sKS(*X* | *Y*) to express the sKS statistics for a pair of distributions of *X* (both continuous in the case of p-STAT1 or both binary otherwise), one obtained for *Y*-negative cells and the other obtained for *Y*-positive cells. The sKS statistic attains the values of ±1 for fully disjointed distributions and 0 for exactly overlapping distributions.

#### Model justification and choice of parameters

The model structure is justified and model parameters are constrained by a series of experiments involving IFNβ stimulation and RSV infection of A549 cells. In Appendix Figs S1–S5, overviewed below, we compare population-averaged model trajectories with respective population-based data.

##### Cascade initiated by RSV infection

In Appendix Fig S1, based on Western blotting and ELISA, we quantified levels of proteins in the cascade initiated by RSV infection at three MOIs (0.01, 0.1 and 1); in Appendix Fig S2, for MOI = 0.1, we analyzed how this cascade is affected by IRF3 knockout. In brief: RSV infection leads to phosphorylation of IRF3 (Appendix Fig S1A), synthesis and secretion of IFNβ (Fig S1C) that in turn activates STAT1 and STAT2 (jointly represented by accumulation of `pSTATs` in the model) and finally accumulation of three proteins coded by interferon stimulated genes: RIG-I, PKR and OAS1 (`ISGs` in the model), as well as STAT1 and STAT2. In IRF3 deficient cells, we observed no IFNβ production (Appendix Fig S1C) and consequently, no STAT1 and STAT2 phosphorylation and no accumulation of ISGs.

##### Dynamics of STATs and ISGs

The IFNβ signaling is key to build antiviral state in bystander cells, and thus we characterized in detail responses to IFNβ in separate experiments. Parameters governing `pSTATs` kinetics were estimated based on Appendix Fig S3A and C. In Appendix Fig S3A, we found that in a broad range of IFNβ stimulation doses (30 U/ml–1000 U/ml) STAT1 phosphorylation peaks and STAT2 phosphorylation plateaus 0.5 hour after stimulation. STAT2 phosphorylation monotonically increases with stimulation dose, reaching half of the maximal activation for about 100 U/ml, and this value determines the Michaelis–Menten constant in `pSTATs` heterodimers accumulation. Appendix Fig S3C shows that one hour after IFNβ washout STAT1 and STAT2 lose phosphorylation, but can be rapidly re-phosphorylated in response to the next IFNβ pulse (which allows estimation of forward and reverse kinetic rate constants for `pSTATs` accumulation).

The forward and reverse kinetic rate constants for ISGs were estimated based on Appendix Fig S3E, in which we observe accumulation of RIG-I, PKR and OAS1 over 24 hours of IFNβ stimulation and their slow degradation over 48 hours after IFNβ washout and STATs dephosphorylation.

Comparison of experimental p-STAT1 and p-STAT2 time profiles with `pSTATs` following from the model (Appendix Fig S3B and D) indicates that the kinetic of p-STAT1 is more complex, but the assumed simplified kinetic of `pSTATs` which follows kinetic of p-STAT2 is sufficient to reproduce kinetics of ISGs (Appendix Fig S3F).

##### Antiviral effect of IFN signaling

In Appendix Fig S4A we studied the importance of IFNβ and IFNλ for attenuating virus spread by knockouts of their respective receptors, IFNAR1 and IFNLR1. We observed that at 48 h p.i. there is significantly more RSV protein F in IFNAR1 KO cells than in WT cells. In turn, IFNLR1 knockout does not influence RSV protein accumulation, as indicated by a comparison of WT and IFNLR1 KO cells as well as of IFNAR1 KO cells and DKO cells (knockout of both IFNAR1 and IFNLR1). This indicates that IFNβ is critical for attenuating RSV spread, and accordingly in our RSV model the ‘generic interferon’ can be identified with IFNβ.

IFNβ attenuates RSV spread by activating STAT1 and STAT2 that dimerize and trigger expression of ISGs. Accordingly, both in STAT1 KO and STAT2 KO cells we did not observe accumulation of ISGs: RIG-I, PKR and OAS1, and observed increased accumulation of RSV protein F (Appendix Fig S4C). Stimulation of WT cells for 24 hours before infection with IFNβ results in a decrease of RSV protein F with respect to non pre-stimulated cells, but this effect is negligible in STAT1 KO and STAT2 KO cells.

The effect of STAT1, STAT2 and interferons receptors knockouts on RSV propagation is reproduced by the model, as shown in Appendix Fig S4B and D. When comparing model with experimental data from Appendix Fig S4A and C, we simulated IFNAR1 KO, DKO (with IFNAR1 and IFNLR1 knockouts), STAT1 KO and STAT2 KO cell by prohibiting `ISGs` from increasing (as in this case the p-STAT1/p-STAT2 dimers cannot be formed), while IFNLR1 KO is treated as WT (i.e., simulated with default parameters). Trajectories p-STAT1 in STAT2 KO and p-STAT2 in STAT1 KO cells are assumed to be the same as in WT cells (as activation of STATs isoforms is independent), but obviously we assumed no increase of p-STAT1 in STAT1 KO cells and no increase of p-STAT2 in STAT2 KO cells.

The inhibitory effect of IFNβ prestimulation on RSV propagation is shown using Western blotting (Appendix Fig S5A) and dPCR (Appendix Fig S5C). Generally, the influence of prestimulation is more pronounced for lower MOIs (where the level of RSV RNA and proteins depends on replication cycles inhibited by ISGs accumulated due to IFNβ prestimulation). This tendency is reproduced by our model (Appendix Fig S5B and C).

#### Model implementation

Agents (cells) are arranged on a triangular lattice (in which each node has 6 neighbors), which mimics a monolayer of epithelial cells (in which, based on Voronoi tessellation with geometric centers of cell nuclei used as seeds, each cell has on average 6 neighboring cells). Periodic boundary conditions are assumed along both axes. In all simulations, 100% confluency is assumed, meaning that all lattice nodes are occupied by cells.

For three variables: `vRNA`, `pIRF3`, and `IFNi` the last state, `3`, stands for active species, while the intermediate states {`1`, `2`} are introduced to account for time delay associated with activation or synthesis. For the remaining three intracellular species: `vProteins`, `pSTATs`, `ISGs`, the intermediate states {`1`, `2`} stand for partial activation (which reflects the assumption of gradual accumulation or activation of these species). Of note, `vProteins = 1`, `2`, `3` may inhibit the increase of `IRF3` and `IFNi`, but only `vProteins = 3` renders the cell infectious.

The stochastic internal state of cells is coupled to an external, two-layer interferon field (see Fig 1B). The field evolves following the diffusion equation, solved using discretization introduced by cell-associated subvolumes. Cells with `IFNi = 3` secrete interferon to the lower layer, and `pSTATs` is activated following Michaelis–Menten kinetics, based on the amount of interferon in this layer. The upper layer accounts for the fact that in experimental settings interferon diffusion takes place in 3D (not 2D) space.

Unless specified otherwise, the model is initialized with all stages and interferon layers set to 0. Infected cells are selected randomly with probability dictated by MOI and their `v` is set to 1. Prestimulation with IFN sets the upper layer to the specified concentration; a wash sets both layers to 0. A typical simulation (100 × 100 grid, 100% cell confluence, IFN field update every 0.1 min) with default parameters and MOI 0.1 for 24 hours of simulated time takes a few seconds of CPU time on a standard PC.

The model does not account for cell movement, and only infection of neighboring cells is allowed. This is consistent with microscopy images where we observe growing clusters of infected cells (see Fig 2). The model also does not account for apoptosis of infected cells, as it is known that apoptosis is blocked by RSV (Thomas *et al*, 2002).

To report population-averaged variables, 1000 stochastic simulations were performed on the 100 × 100 grid, with the exception of Figures 3D, 7 and Appendix Figures S1–S5, S7, S8A for which 100 stochastic simulations were performed. The estimations of credible intervals (Fig. 3C) was based on 10,000 stochastic simulations.

#### Comparison of experimental and simulation results

MOI and the time post infection are two parameters that in principle should determine the state of infection. Because of unavoidable differences between experimental replicates in the number of active virus particles actually infecting cells (despite the same MOI) we observe that using the observed fraction of cells expressing RSV proteins in a given time post infection (instead of MOI) allows for more reliable comparisons across experiments, and with model simulations.

Neighboring cells are well defined for simulations (on the model lattice each cell has 6 neighbors). In order to compare simulation results with experimental data, we define neighbors by constructing Voronoi diagrams for microscope images. We identify cell nuclei using DAPI staining and use their centers of mass as Voronoi centers. We then consider cells to be neighbors if their respective Voronoi tiles share an edge. Conveniently, this procedure assigns on average 6 neighbors to each cell, the same as in our model.

In immunostaining images, levels of observed species IFNβ, nuclear IRF3 and RSV proteins are discretized into binary variables (quantification and discretization of immunostaining images is described in previous subsection Image quantification). We perform a similar binarization in the model when comparing with immunostaining images. In the case of `IFNi` and `pIRF3`, for which the states {`1`, `2`} are introduced solely to account for production delay, only their last state, `3` is considered active. In the case of `vProteins` intermediate states denote accumulation of viral proteins, and thus all states `vProteins > 0` are considered active. In analysis of experimental data we leave p-STAT1 as continuous variable, while in the model we use 4-state discretization. Importantly, this means that for low levels of p-STAT1, small differences might be picked up in experimental data but are not accounted for in the model.

To compare Western blots with the model simulation data, for a given simulation protocol we performed 100 simulations on a 100 × 100 lattice, and then average over all cells and simulations. We assumed that the levels of p-STAT1 and p-STAT2 are proportional to the average activation level for `pSTATs`, the levels of RIG-I, PKR and OAS1 are proportional to the average activation level of `ISGs`, and the level of RSV protein F is proportional to the average activation level of `vProteins.` We also assumed that the level of p-IRF3 is proportional to the fraction of cells in which `pIRF3 = 3` (as in this case state `1` and `2` are introduced solely to account for time delay associated with IRF3 activation). Western blots were first normalized by the corresponding reference GAPDH, then by the maximum value in the series of measurements (from the same blot). Next, we added 0.03 to each value and finally normalized the series by the geometric mean. This procedure reflects the assumption that blots adequately capture fold differences between measurements and that differences between values smaller than 0.03 of the maximum from a given blot are not meaningful. The same normalization procedure (without the reference GAPDH step) is applied to simulation data. Overall, this approach yields a dynamic range that accommodates up to ∼30-fold protein level differences.

ELISA measurements (Appendix Fig S1C) are compared with `IFNe` summed over upper and lower compartments and averaged over cells. dPCR measurements for RSV RNA (Appendix Fig S5C) are compared with the model based on the simplifying assumption that the average level of viral RNA is proportional to the fraction of cells with `vRNA = 3` (for `vRNA` states `1` and `2` account solely for time delay).

## Acknowledgements

This study was funded by the National Science Centre Poland grants 2019/34/H/NZ6/00699 (Norwegian Financial Mechanism) and 2018/29/B/NZ2/00668.

## Data Availability

Source code of the agent-based simulator of the innate immune response to an infection with an RNA virus is available at https://github.com/grfrederic/visavis. The simulator implements the computational model described in and used throughout the article.

## Appendix

Note

In Appendix Figs S1, S3, S4, and S5 we juxtapose model predictions with quantified experimental results reported in:

- Czerkies M, Kochańczyk M, Korwek Z, Prus W & Lipniacki T (2022) Respiratory syncytial virus protects bystander cells against influenza A virus infection by triggering secretion of type I and type III Interferons. *J Virol*: e0134122 10.1128/jvi.01341-22.
- Korwek Z, Czerkies M, Jaruszewicz-Błońska J, Prus W, Kosiuk I, Kochańczyk M & Lipniacki T (2022) Non-self RNA rewires IFNβ signaling: A mathematical model of the innate immune response. *bioRxiv* doi:10.1101/2022.01.30.478391 (preprint).

as detailed in figure legends. For the reader’s convenience, we show corresponding Western blot replicates (not shown in these papers).

**Appendix Figure S1.**
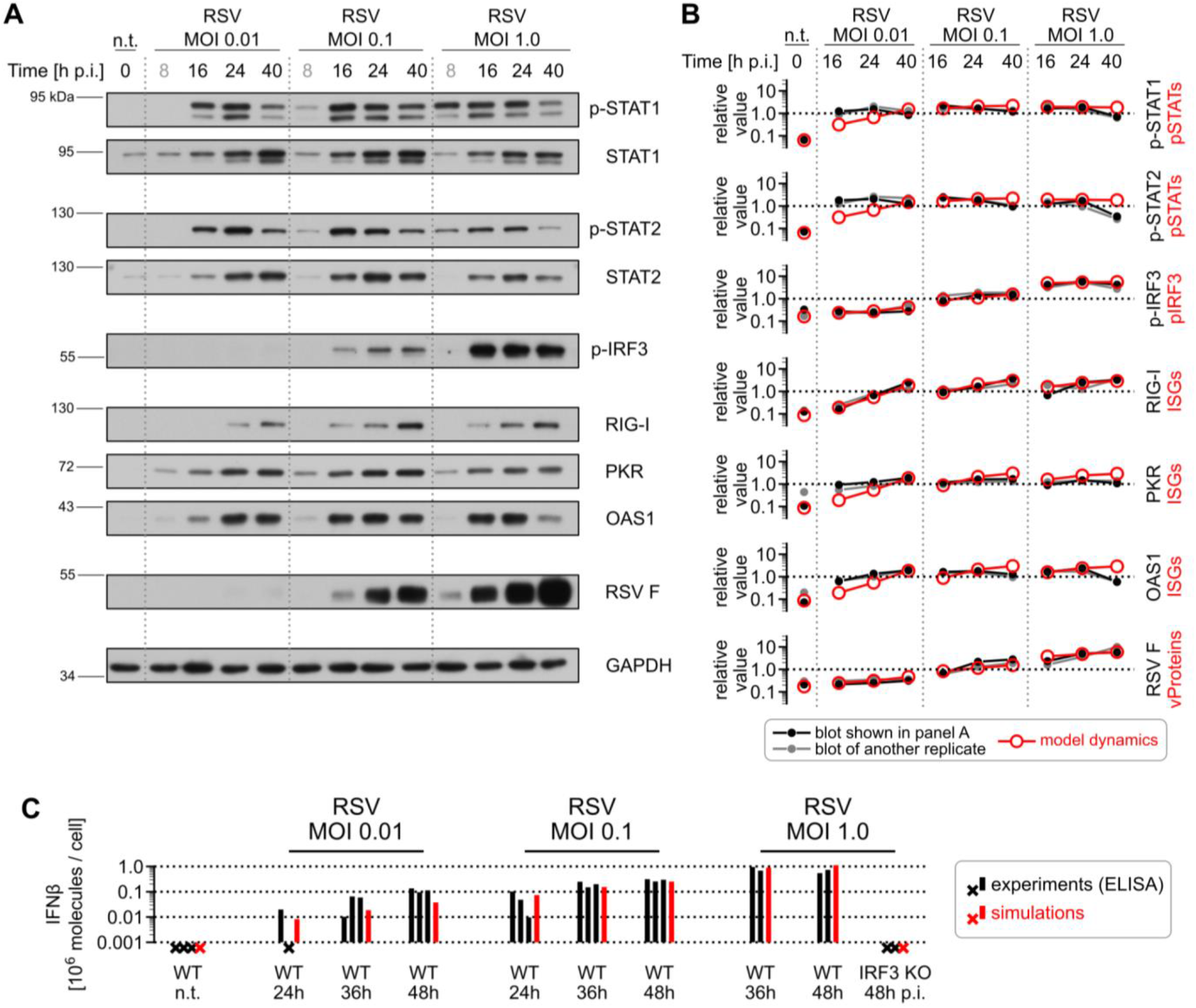
Progression of RSV infection without IFNβ pre-stimulation. **A, B** Progression of RSV infection at MOIs of 0.01, 0.1, and 1 in A549 cells. Western blot analysis (A, B) is juxtaposed with model simulations (B). Time point of 8 h p.i. lacks experimental replicates and is thus not included in panel C. **C** Infected cells produce and secrete IFNβ. Results of ELISA for A549 cells infected with RSV at MOIs of 0.01, 0.1, and 1 (black) are juxtaposed with model predictions (red). Experiments with samples below the limit of detection as well as simulation results with no IFN are marked with crosses. For MOIs of 0.01 and 0.1 we used data from Czerkies *et al* (2022, see Figure 1A therein).

**Appendix Figure S2.**
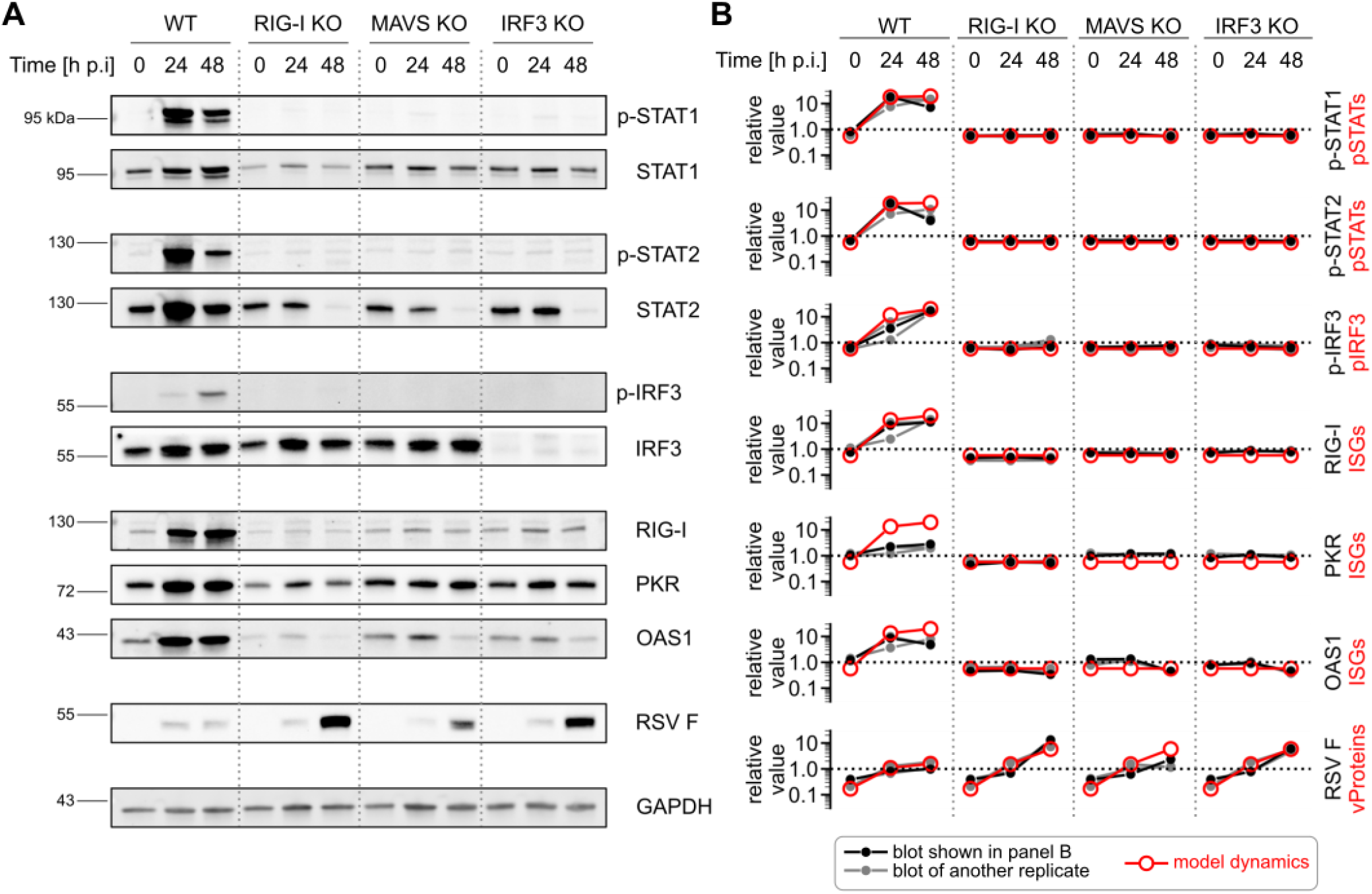
Viral infection triggers an innate immune signaling cascade. **A, B** RSV activates multiple components of the innate immune signaling. Western blot analysis of A549 cells infected with RSV at an MOI of 0.1 (A, B) is juxtaposed with model simulations (B). In panel B, black circles and connecting lines are quantifications of blots shown in panel A, gray filled circles and connecting lines are quantifications of blots from another experimental replicate, and red circles and connecting lines were obtained by sampling the average model trajectory at experimental time points. All KO cell lines were simulated by disabling activation of `pIRF3`.

**Appendix Figure S3.**
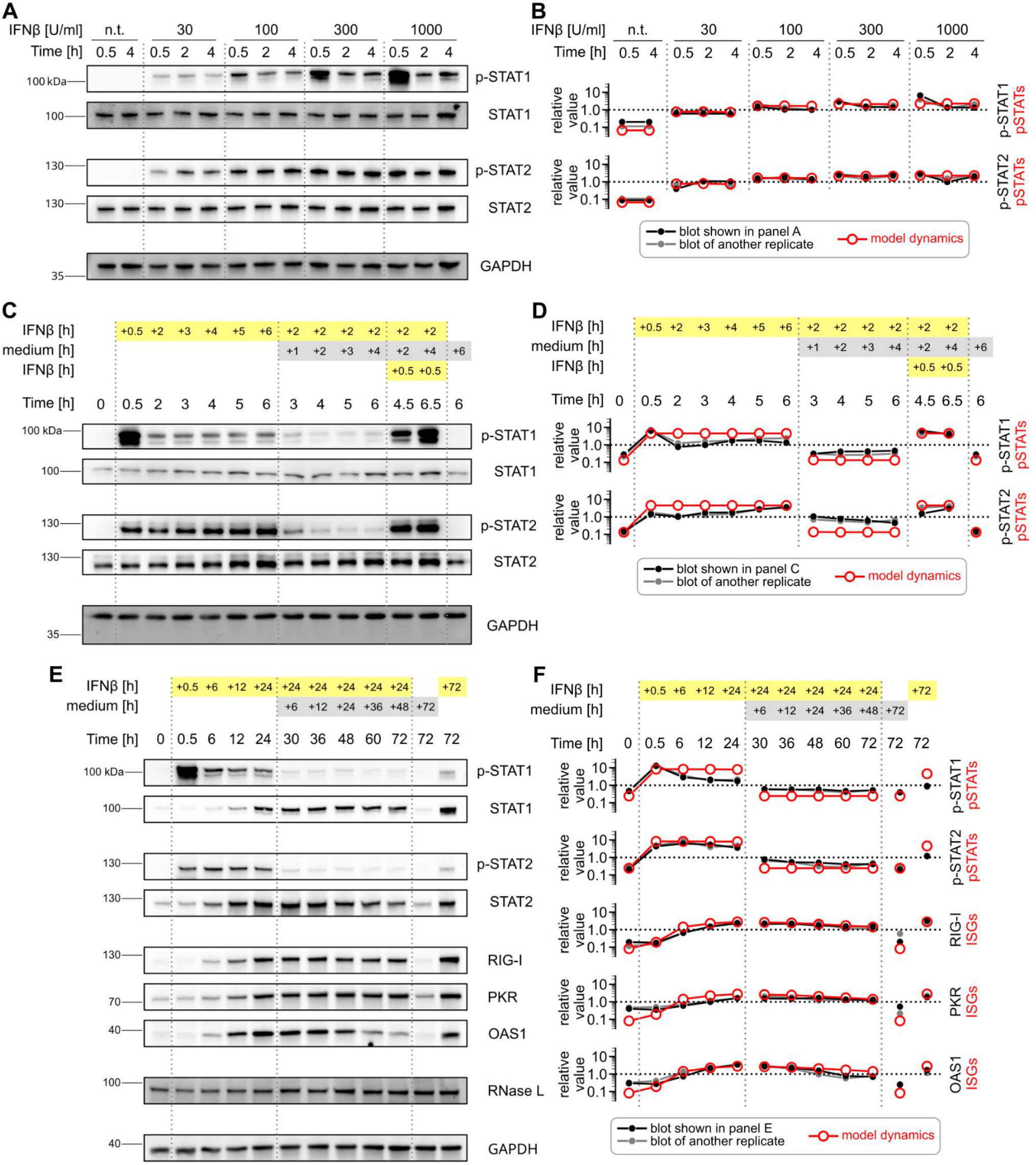
STAT1/2 respond quickly but IFNβ-stimulated proteins accumulate and degrade slowly. **A, B** STAT1/2 are activated quickly and their amplitude depends on IFNβ concentration up to 1000 U/ml. Western blot analysis of A549 cells (A, B) is juxtaposed with model simulations (B). Western blot data originates from Korwek *et al* (2022; see Extended Data Fig. 1c therein). The comparison between model and experiment in panel B is based on the conversion 1000 U/ml = 5.5 ng/ml for the IFNβ stock used. **C, D** IFNβ is required to sustain STAT1/2 activity. A549 cells were stimulated with IFNβ (1000 U/ml) according to the indicated protocols: for example, in the next to last column, cells were stimulated with IFNβ for 2 h, then IFNβ was washed out by medium replacement and cells were cultured in a fresh medium for subsequent 4 h, and finally IFNβ was added for 0.5 h (protocol total duration is 6.5 h). Western blot analysis (C, D) is juxtaposed with model simulations (D). Western blot data originates from Korwek *et al* (2022; see Fig. 3a therein). **E, F** IFNβ-stimulated proteins accumulate and degrade slowly. A549 WT cells were stimulated with IFNβ (1000 U/ml) according to the indicated protocols. Western blot analysis (E, F) is juxtaposed with model simulations (F).

**Appendix Figure S4.**
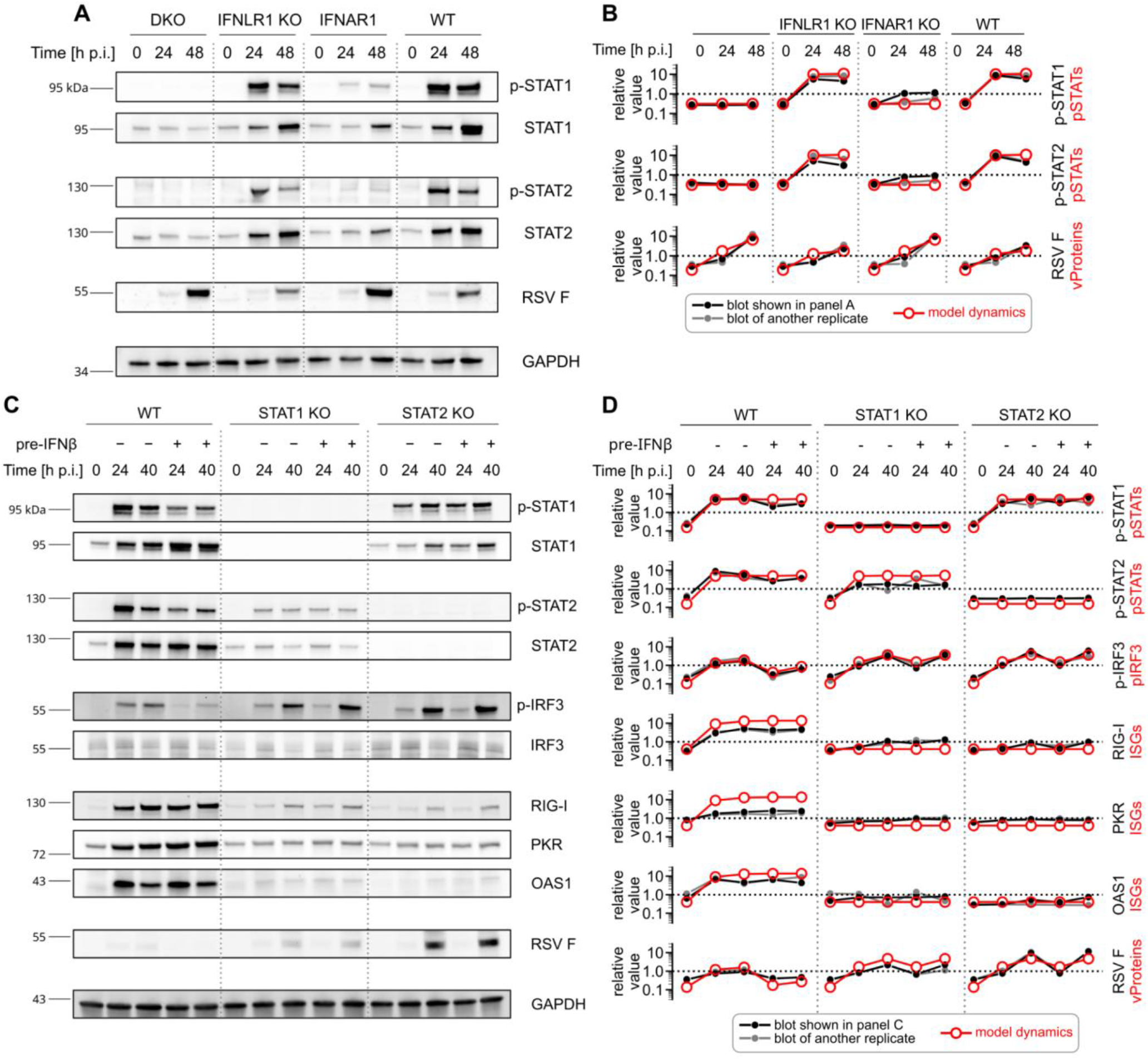
IFNβ/λ-induced STAT1/2 signaling attenuates viral infection. **A, B** Both IFNβ and, to a lesser degree, IFNλ activate STAT1 and STAT2. Western blot analysis of A549 cells infected with RSV at an MOI of 0.1 (A, B) is juxtaposed with model simulations (B). Western blot data originates from Czerkies *et al* (2022; see Figure 1D therein). Both the IFNAR1 KO and IFNAR1–INFLR1 double KO cell lines were simulated by disabling the forward transition of `pSTATs`. **C, D** Activation of STAT1/2 attenuates the spread of RSV infection. Western blot analysis of A549 cells infected with RSV at an MOI of 0.1 (C, D) is juxtaposed with model simulations (D). Both the STAT1 KO and STAT2 KO cell lines were simulated by disabling the forward transition of `pSTATs`.

**Appendix Figure S5.**
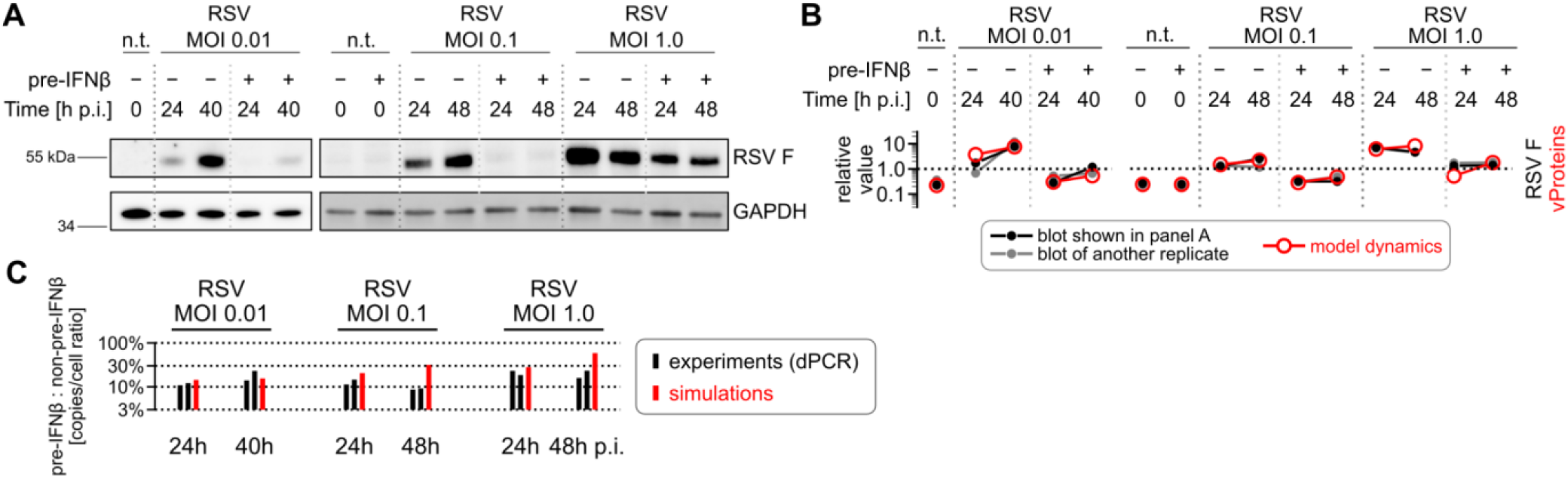
Pre-stimulation with IFNβ impedes virus spread. **A, B** Propagation of RSV infection and signaling without or with prior IFNβ pre-stimulation (1000 U/ml). A549 cells were infected with RSV at MOI of 0.01, 0.1, and 1. Western blot analysis (A, B) is juxtaposed with model simulations (B). For MOIs 0.1, 1 we use experimental data shown in Czerkies *et al* (2022; as a part of Figure 2C therein). **C** Viral load in RSV infection without or with prior IFNβ pre-stimulation (1000 U/ml). Digital PCR results for A549 cells infected with RSV at MOIs of 0.01, 0.1, and 1 (black) are juxtaposed with model predictions (red). For an MOI of 0.1 and 1 we use data shown in Czerkies *et al* (2022; see Figure 2A therein).

**Appendix Figure S6.**
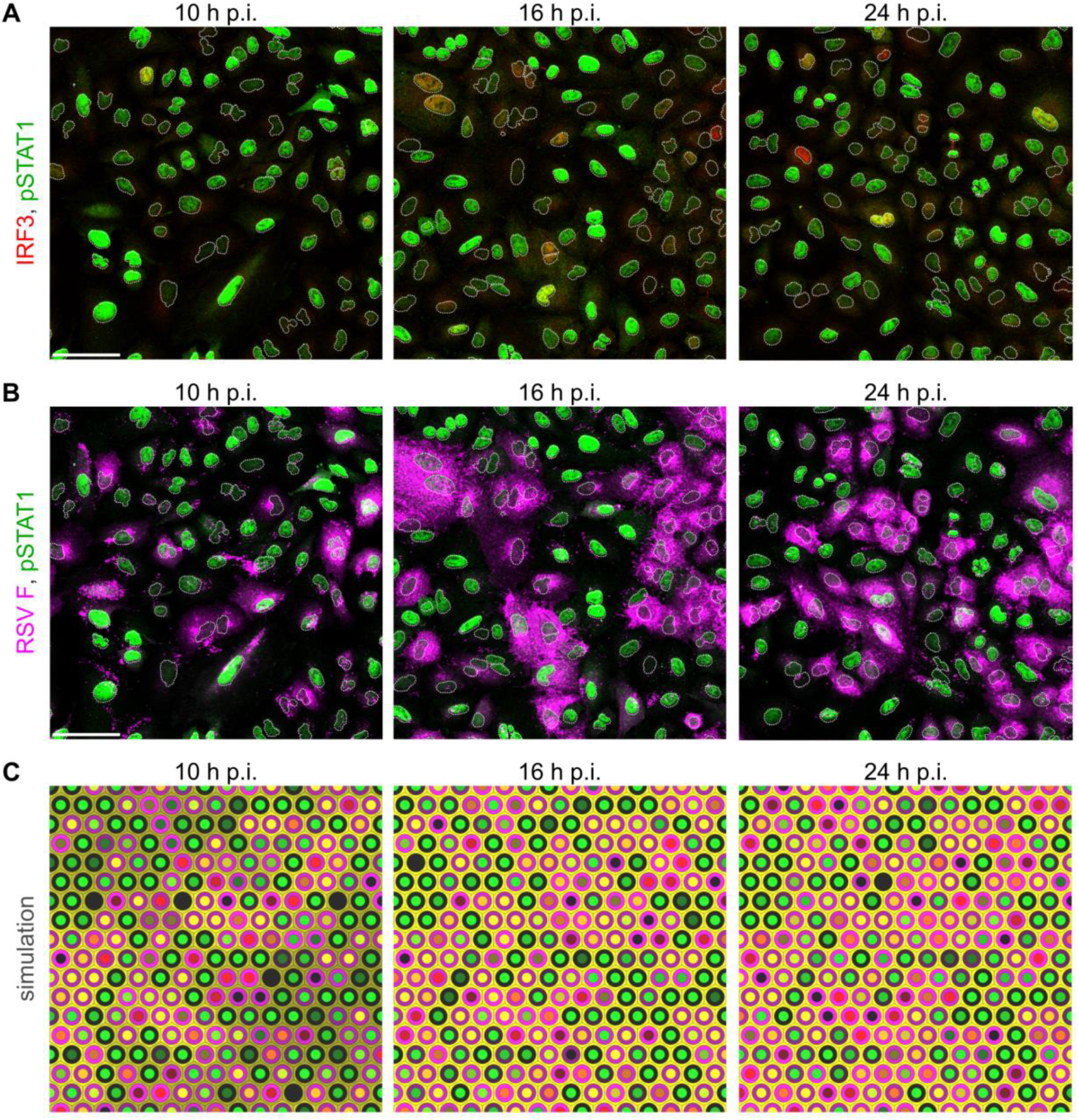
Images from experiment and snapshots from simulation, infection at an MOI of 1. **A, B** A549 cells 10, 16, and 24 hours post infection (p.i.) with RSV at an MOI of 1. IRF3 – red (intracellular, only in panel A), RSV – magenta (only in panel B), p-STAT1 – green. White dotted lines are nuclear outlines determined based on DAPI counterstaining (channel not shown). Scale bars, 50 µm. **C** Snapshots from a simulation of infection at an MOI of 1 in a compact monolayer of cells (subpanels show small fragments of a simulated 100×100 lattice).

**Appendix Figure S7.**
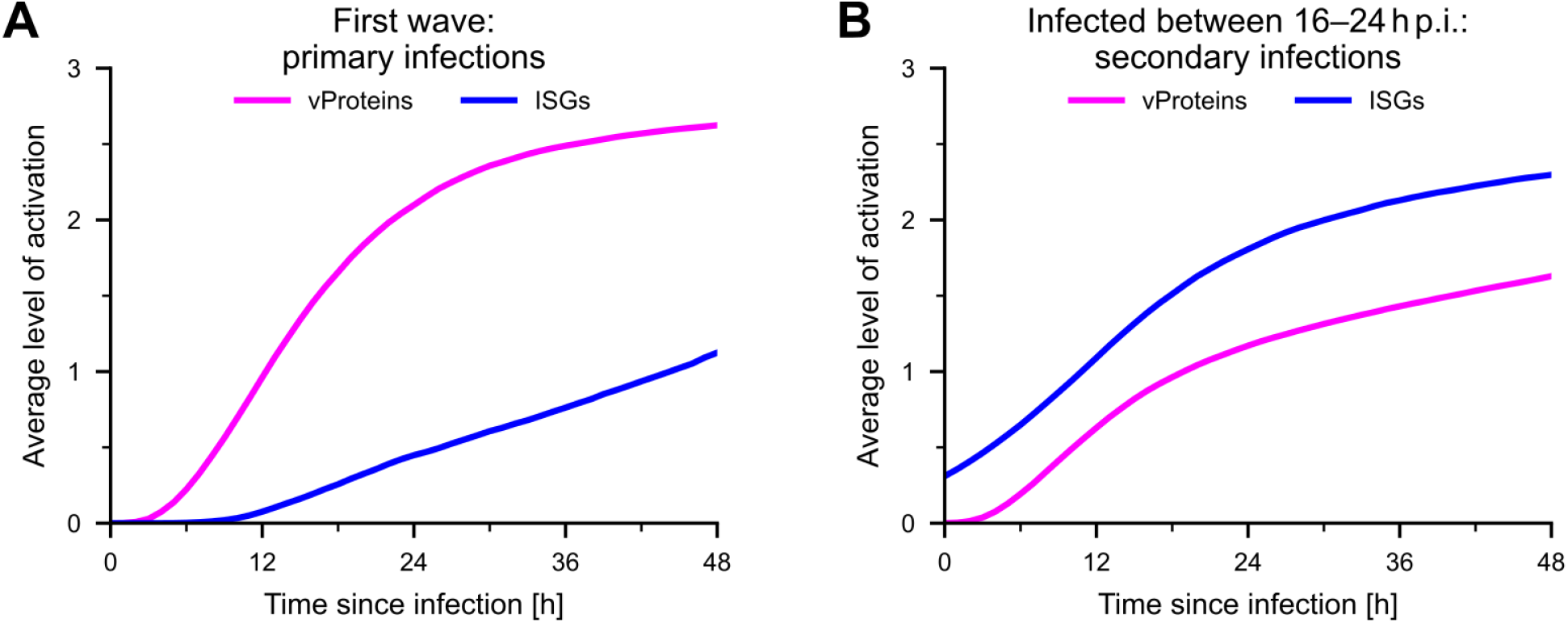
Viral proteins accumulate more slowly in cells protected by ISGs (model). **A, B** Average status of `vProteins` and `ISGs` as a function of time since infection of individual cells. A simulated cell culture was infected at an MOI of 0.01. Panel A shows primarily infected cells, whereas panel B shows cells that were infected in between 16 and 24 h post infection of the simulated cell culture (secondary infections). In panel A, the time axis shows absolute time post infection; in panel B, time axes of individual cells were aligned to begin at their particular time of infection.

**Appendix Figure S8.**
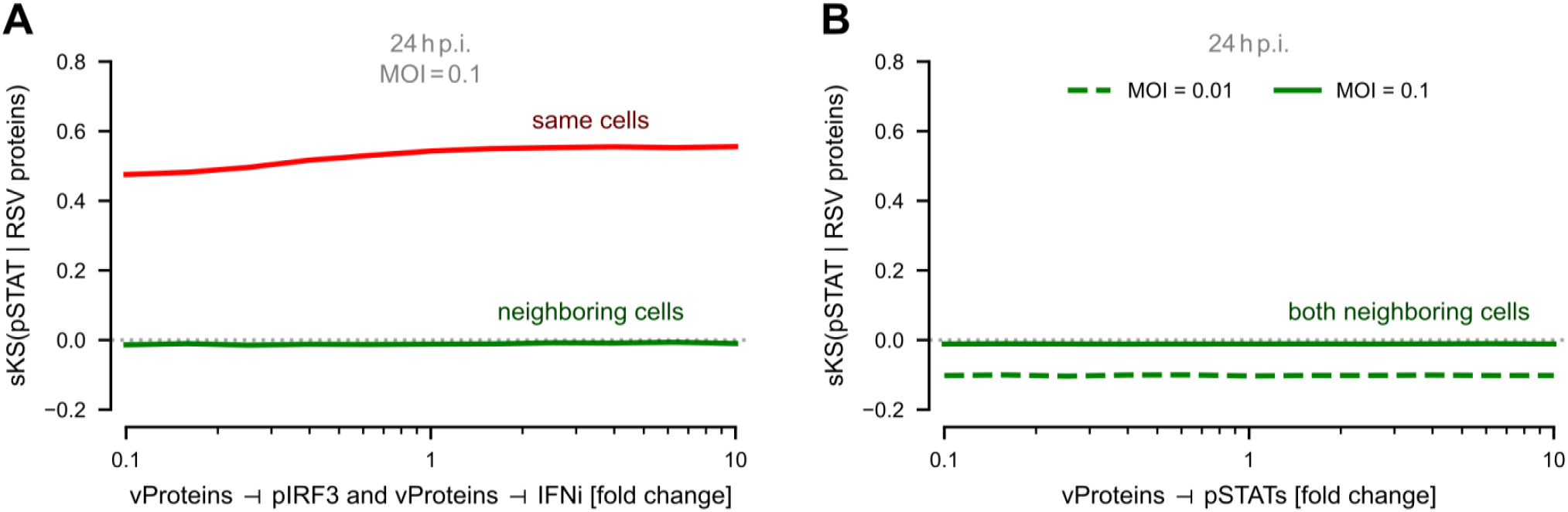
Lack of dependence of the signed Kolmogorov–Smirnov statistics for (pSTAT | RSV proteins) on remaining inhibition strengths (model). **A** ‘Same cells’ and ‘neighboring cells’ sKS statistics at 24 h p.i. and MOI = 0.1 as a function of the strengths of `vProteins` ⊣ `pIRF3` and `vProteins` ⊣ `IFNi` inhibitions (both strengths were varied simultaneously). **B** ‘Neighboring cells’ sKS statistics at 24 h p.i., for two MOIs (0.01, 0.1) as a function of the strength of the `vProteins` ⊣ `pSTATs` inhibition.

**Appendix Table S1.** Model equations and parameters.

| Process | Rate | Parameter values |
| --- | --- | --- |
| (1) infection | $[v = 0] \times v\_incr$<br>$\times (\# \text{ of neighbors with } vProteins = 3)$ | $v\_incr = 0.25 / h$ |
| (2) vRNA forward | $[vRNA < 3] \times v \times vrna\_incr$<br>$/ (1 + isg\_inh\_vrna \times ISGs)$ | $vrna\_incr = 0.5 / h$<br>$isg\_inh\_vrna = 2.0$ |
| (3) vProteins forward | $[vProteins < 3] \times [vRNA = 3] \times vprot\_incr$<br>$/ (1 + isg\_inh\_vprot \times ISGs)$ | $vprot\_incr = 0.167 / h$<br>$isg\_inh\_vprot = 2.0$ |
| (4) pIRF3 forward | $[pIRF3 < 3] \times [vRNA = 3] \times pirf3\_incr$<br>$/ (1 + vprot\_inh\_pirf3 \times vProteins)$ | $pirf3\_incr = 0.75 / h$<br>$vprot\_inh\_pirf3 = 3.0$ |
| (5) pIRF3 backward | $[pIRF3 > 0] \times pirf3\_decr$ | $pirf3\_decr = 0.125 / h$ |
| (6) IFNi forward | $[IFNi < 3] \times [pIRF3 = 3] \times ifni\_incr$<br>$/ (1 + vprot\_inh\_ifni \times vProteins)$ | $ifni\_incr = 1.0 / h$<br>$vprot\_inh\_ifni = 2$ |
| (7) IFNi backward | $[IFNi > 0] \times ifni\_decr$ | $ifni\_decr = 0.25 / h$ |
| (8) increase of IFNe | $[IFNi = 3] \times k\_ifn\_sec$ | $ifn\_sec = 5e5 / h$ |
| (9) decrease of IFNe | $q\_ifne$ | $q\_ifne = 1 / \text{day}$ |
| (10) pSTATs forward | $[pSTATs < 3] \times pstat\_incr$<br>$\times IFNe / (mm\_pstat + IFNe)$<br>$/ (1 + vprot\_inh\_pstat \times vProteins)$ | $pstat\_incr = 40.0 / h$<br>$mm\_pstat = 500$<br>$vprot\_inh\_pstat = 1.5$ |
| (11) pSTATs backward | $[pSTATs > 0] \times pstat\_decr$ | $pstat\_decr = 10.0 / h$ |
| (12) ISGs forward | $[ISGs < 3] \times isg\_incr \times (pSTATs / 3)$ | $isg\_incr = 0.3 / h$ |
| (13) ISGs backward | $[ISGs > 0] \times isg\_decr$ | $isg\_decr = 0.033 / h$ |

In rate expressions, $[condition]$ is defined to be 1 if *condition* is true and 0 otherwise (Iverson bracket).

## References

Aponte-Serrano JO, Weaver JJA, Sego TJ, Glazier JA & Shoemaker JE (2021) Multicellular spatial model of RNA virus replication and interferon responses reveals factors controlling plaque growth dynamics. PLoS Comput Biol 17: e1008874

Ayllon J & García-Sastre A (2015) The NS1 protein: a multitasking virulence factor. Curr Top Microbiol Immunol 386: 73–107

Barber GN (2001) Host defense, viruses and apoptosis. Cell Death Differ 8: 113–126

Barik S (2013) Respiratory syncytial virus mechanisms to interfere with type 1 interferons. Curr Top Microbiol Immunol 372: 173–191

Bitko V, Shulyayeva O, Mazumder B, Musiyenko A, Ramaswamy M, Look DC & Barik S (2007) Nonstructural proteins of respiratory syncytial virus suppress premature apoptosis by an NF-kappaB-dependent, interferon-independent mechanism and facilitate virus growth. J Virol 81: 1786–1795

Crowe JE (2014) Human Respiratory Viruses. Ref Module Biomed Sci: B978-0-12-801238-3.02600-3

Czerkies M, Kochańczyk M, Korwek Z, Prus W & Lipniacki T (2022) Respiratory syncytial virus protects bystander cells against influenza A virus infection by triggering secretion of type I and type III Interferons. J Virol: e0134122

Czerkies M, Korwek Z, Prus W, Kochańczyk M, Jaruszewicz-Błońska J, Tudelska K, Błoński S, Kimmel M, Brasier AR & Lipniacki T (2018) Cell fate in antiviral response arises in the crosstalk of IRF, NF-κB and JAK/STAT pathways. Nat Commun 9: 493

Drayman N, Patel P, Vistain L & Tay S (2019) HSV-1 single-cell analysis reveals the activation of anti-viral and developmental programs in distinct sub-populations. eLife 8: e46339

Falsey AR, Hennessey PA, Formica MA, Cox C & Walsh EE (2005) Respiratory syncytial virus infection in elderly and high-risk adults. N Engl J Med 352: 1749–1759

García-Sastre A (2017) Ten Strategies of Interferon Evasion by Viruses. Cell Host Microbe 22: 176–184

Gregg RW, Shabnam F & Shoemaker JE (2021) Agent-based modeling reveals benefits of heterogeneous and stochastic cell populations during cGAS-mediated IFNβ production. Bioinformatics 37: 1428–1434

Hale BG, Albrecht RA & García-Sastre A (2010) Innate immune evasion strategies of influenza viruses. Future Microbiol 5: 23–41

Korwek Z, Czerkies M, Jaruszewicz-Błońska J, Prus W, Kosiuk I, Kochańczyk M & Lipniacki T (2022) Non-self RNA rewires IFNβ signaling: A mathematical model of the innate immune response. 2022.01.30.478391 doi:10.1101/2022.01.30.478391[PREPRINT]

Maier BD, Aguilera LU, Sahle S, Mutz P, Kalra P, Dächert C, Bartenschlager R, Binder M & Kummer U (2022) Stochastic dynamics of Type-I interferon responses. PLoS Comput Biol 18: e1010623

Patil S, Fribourg M, Ge Y, Batish M, Tyagi S, Hayot F & Sealfon SC (2015) Single-cell analysis shows that paracrine signaling by first responder cells shapes the interferon-β response to viral infection. Sci Signal 8: ra16–ra16

Rand U, Rinas M, Schwerk J, Nöhren G, Linnes M, Kröger A, Flossdorf M, Kály-Kullai K, Hauser H, Höfer T, et al (2012) Multi-layered stochasticity and paracrine signal propagation shape the type-I interferon response. Mol Syst Biol 8: 584–584

Rashid F, Xie Z, Suleman M, Shah A, Khan S & Luo S (2022) Roles and functions of SARS-CoV-2 proteins in host immune evasion. Front Immunol 13: 940756

Sa Ribero M, Jouvenet N, Dreux M & Nisole S (2020) Interplay between SARS-CoV-2 and the type I interferon response. PLOS Pathog 16: e1008737

Scheltema NM, Gentile A, Lucion F, Nokes DJ, Munywoki PK, Madhi SA, Groome MJ, Cohen C, Moyes J, Thorburn K, et al (2017) Global respiratory syncytial virus-associated mortality in young children (RSV GOLD): a retrospective case series. Lancet Glob Health 5: e984–e991

Schoggins JW (2019) Interferon-Stimulated Genes: What Do They All Do? Annu Rev Virol 6: 567–584

Sedeyn K, Schepens B & Saelens X (2019) Respiratory syncytial virus nonstructural proteins 1 and 2: Exceptional disrupters of innate immune responses. PLoS Pathog 15: e1007984

Sego TJ, Aponte-Serrano JO, Gianlupi JF & Glazier JA (2021) Generation of multicellular spatiotemporal models of population dynamics from ordinary differential equations, with applications in viral infection. BMC Biol 19: 196

Sego TJ, Mochan ED, Ermentrout GB & Glazier JA (2022) A multiscale multicellular spatiotemporal model of local influenza infection and immune response. J Theor Biol 532: 110918

Sen A, Rothenberg ME, Mukherjee G, Feng N, Kalisky T, Nair N, Johnstone IM, Clarke MF & Greenberg HB (2012) Innate immune response to homologous rotavirus infection in the small intestinal villous epithelium at single-cell resolution. Proc Natl Acad Sci U S A 109: 20667–20672

Sun Y, Jain D, Koziol-White CJ, Genoyer E, Gilbert M, Tapia K, Panettieri RA, Hodinka RL & López CB (2015) Immunostimulatory Defective Viral Genomes from Respiratory Syncytial Virus Promote a Strong Innate Antiviral Response during Infection in Mice and Humans. PLoS Pathog 11: e1005122

Talemi SR & Höfer T (2018) Antiviral interferon response at single-cell resolution. Immunol Rev 285: 72–80

Thomas KW, Monick MM, Staber JM, Yarovinsky T, Carter AB & Hunninghake GW (2002) Respiratory syncytial virus inhibits apoptosis and induces NF-kappa B activity through a phosphatidylinositol 3-kinase-dependent pathway. J Biol Chem 277: 492–501

Van Eyndhoven LC, Singh A & Tel J (2021) Decoding the dynamics of multilayered stochastic antiviral IFN-I responses. Trends Immunol 42: 824–839

Vignuzzi M & López CB (2019) Defective viral genomes are key drivers of the virus-host interaction. Nat Microbiol 4: 1075–1087

Zhao M, Zhang J, Phatnani H, Scheu S & Maniatis T (2012) Stochastic expression of the interferon-β gene. PLoS Biol 10: e1001249

